# The host environment rather than biofilm growth activates stress responses and conveys ciprofloxacin tolerance in *Staphylococcus aureus*

**DOI:** 10.1101/2025.08.07.669083

**Authors:** Maiken Engelbrecht Petersen, Amanda Batoul Khamas, Cecilie Aaberg Meltofte, Freja Cecilie Mikkelsen, Emma Iben Boskov Hundevadt, Marie Eline Hoppe, Thomas Keith Wood, Rikke Louise Meyer

## Abstract

**Introduction:** Antibiotic tolerance allows genetically susceptible bacteria to survive antibiotic exposure and contributes to treatment failure and recurrent infection. In *Staphylococcus aureus*, tolerance is closely associated with biofilm infections, and this association is conventionally explained by the biofilm mode of growth itself, in which mass transfer limitation is presumed to starve and arrest cells in the deeper layers.

**Gap Statement:** That explanation rests largely on biofilms grown in rich laboratory media, and it has rarely been tested against the alternative that the host environment, rather than the biofilm architecture, drives the underlying physiological change. Furthermore, because transcriptomic and bulk fluorescence methods average across the population, they cannot establish whether stress responses are activated uniformly in all cells or strongly in a minority.

**Aim:** To determine, at single-cell resolution, whether bacterial stress responses are activated by biofilm growth, by the host environment, or by both, and whether such activation coincides with a reduced rate of antibiotic killing.

**Methodology:** Promoter regions of *recA*, *katA*, *relQ*, *ilvleu* and *groESL* were fused to a stable green fluorescent protein variant to report the SOS, oxidative stress, cell wall stress- and starvation-activated stringent, and heat shock responses. Reporter activation was quantified by confocal microscopy in exponential and stationary cultures, after defined chemical and physical stresses, in biofilms grown in tryptic soy broth, in diluted and undiluted human serum, in heat-inactivated serum, and in *S. aureus* internalised by human neutrophils. Tolerance was assessed as the rate of killing in time-kill assays.

**Results:** Biofilms grown in laboratory media or in diluted serum showed minimal stress response activation, whereas biofilms of the same strain grown in undiluted human serum activated all five responses, uniformly across the population. Heat inactivation of the serum did not abolish this effect, indicating that complement and heat-labile serum enzymes are not required. One hour of serum exposure activated the stringent response via both starvation and cell wall stress, and it abolished detectable killing by ciprofloxacin, while producing only small changes in killing by vancomycin and rifampicin and none by daptomycin or dicloxacillin. Pre-treatments that activated the stringent or heat shock responses likewise arrested growth and slowed ciprofloxacin killing. Neutrophil phagocytosis activated the oxidative stress response and both stringent response branches.

**Conclusion:** The host environment, rather than the biofilm mode of growth, is the dominant driver of stress response activation in *S. aureus*. Activation is population-wide and coincides with population-wide ciprofloxacin tolerance. Laboratory biofilm models that omit host components are therefore likely to underestimate the tolerance encountered during infection.

## Introduction

Antibiotic tolerance is a phenotypic trait that allows genetically susceptible bacteria to survive antibiotic exposure, and it is a recognised contributor to treatment failure and recurrent infections ^1^. Tolerance is defined operationally: a tolerant population is killed more slowly than a susceptible one, without any change in the minimum inhibitory concentration ^2^. This phenomenon is commonly associated with bacterial biofilms, making biofilm infections notoriously difficult to treat despite the causative bacteria being susceptible to antibiotics when tested under laboratory conditions.

*Staphylococcus aureus* is among the pathogens most frequently implicated in difficult-to-treat infections, including prosthetic vascular graft infections ^3^, endocarditis, osteomyelitis, and skin and soft tissue infections ^4^. These infections are commonly associated with biofilms, which protect bacteria from the immune system. Biofilms respond poorly to antibiotics, and this phenotype is conventionally attributed to the biofilm mode of growth itself. Mass transfer limitation generates gradients of oxygen and nutrients, and the resulting starvation and growth arrest in the deeper layers are presumed to render the bacteria tolerant to antibiotics because conventional antibiotics are most effective against actively growing bacteria. This model is intuitive and widely applied, but it has largely been built from biofilms grown in rich laboratory media, and it has rarely been tested against the alternative possibility that the host milieu, rather than the biofilm architecture, is what drives the underlying physiological change.

Cellular stress responses are plausible mediators of that change. Tolerance in *S. aureus* has been linked to low metabolic activity ^5^, starvation ^6, 7^, inhibition of transcription, translation or ATP synthesis ^8^, protein phosphorylation ^9^, and to the activation of stress responses including the SOS response ^10–13^, the oxidative stress response ^14–16^, the stringent response ^12, 17–21^ and the heat shock response ^12^. Several of these links have been established under host-relevant conditions. Ledger et al. ^22^ showed that exposure to human serum triggers tolerance to multiple antibiotics. Peyrusson et al. ^12, 15^ showed that intracellular *S. aureus* survives antibiotic exposure and mounts a broad stress response, driven in part by host-derived oxidative stress; Rowe et al. ^16^ showed that reactive oxygen species induce tolerance during systemic infection; and Huemer et al. ^9^ recovered growth-arrested, antibiotic-tolerant bacteria directly from patient material. Mechanistic dissection has proved less straightforward: a mutant unable to synthesise (p)ppGpp remained as susceptible to ciprofloxacin as the wild type, leading to the proposal that tolerance arises from ATP depletion rather than from alarmone signalling as such ^5^.

Two questions are left open by this body of work, and both are consequences of the methods used. First, because serum exposure and biofilm growth have generally been studied separately, it is not established whether the biofilm mode of growth is itself sufficient to activate stress responses, or whether the host environment is the operative variable. Second, because transcriptomic and bulk fluorescence measurements average across the population, they cannot distinguish uniform activation in all cells from strong activation in a minority. That distinction is precisely what separates tolerance from persistence, in which a dormant sub-population ensures survival antibiotic treatment, only to activate and repopulate after treatment has ended. Distinction between population-wide vs sub-population phenomena cannot be resolved without single-cell resolution.

We therefore constructed a set of fluorescent transcriptional reporters that allow stress response activation to be read out in individual *S. aureus* cells, in situ, under conditions that cannot be sampled by bulk methods — within intact biofilms and inside neutrophils. Promoter regions of five genes were fused to GFPuvr, a stable and strongly fluorescent GFP variant ^23^: *recA* (SOS response), *katA* (oxidative stress response), *relQ* and *ilvleu* (the cell wall stress- and starvation-activated branches of the stringent response, respectively), and *groESL* (heat shock response). The rationale for each reporter, the stress it reports, and the conditions used to induce it are given in Table 1. We note at the outset that *groESL* reports the accumulation of misfolded protein rather than elevated temperature specifically, and that it can therefore be activated by conditions unrelated to heat, including oxidative damage and disruption of protein synthesis ^24^. We have made all five reporter strains available through Addgene and they are carried on a shuttle plasmid that can be introduced into clinical isolates.

**Table 1.**
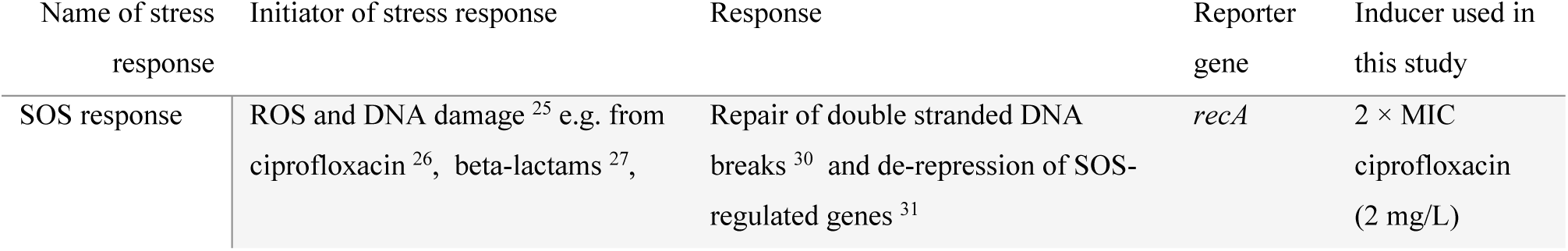
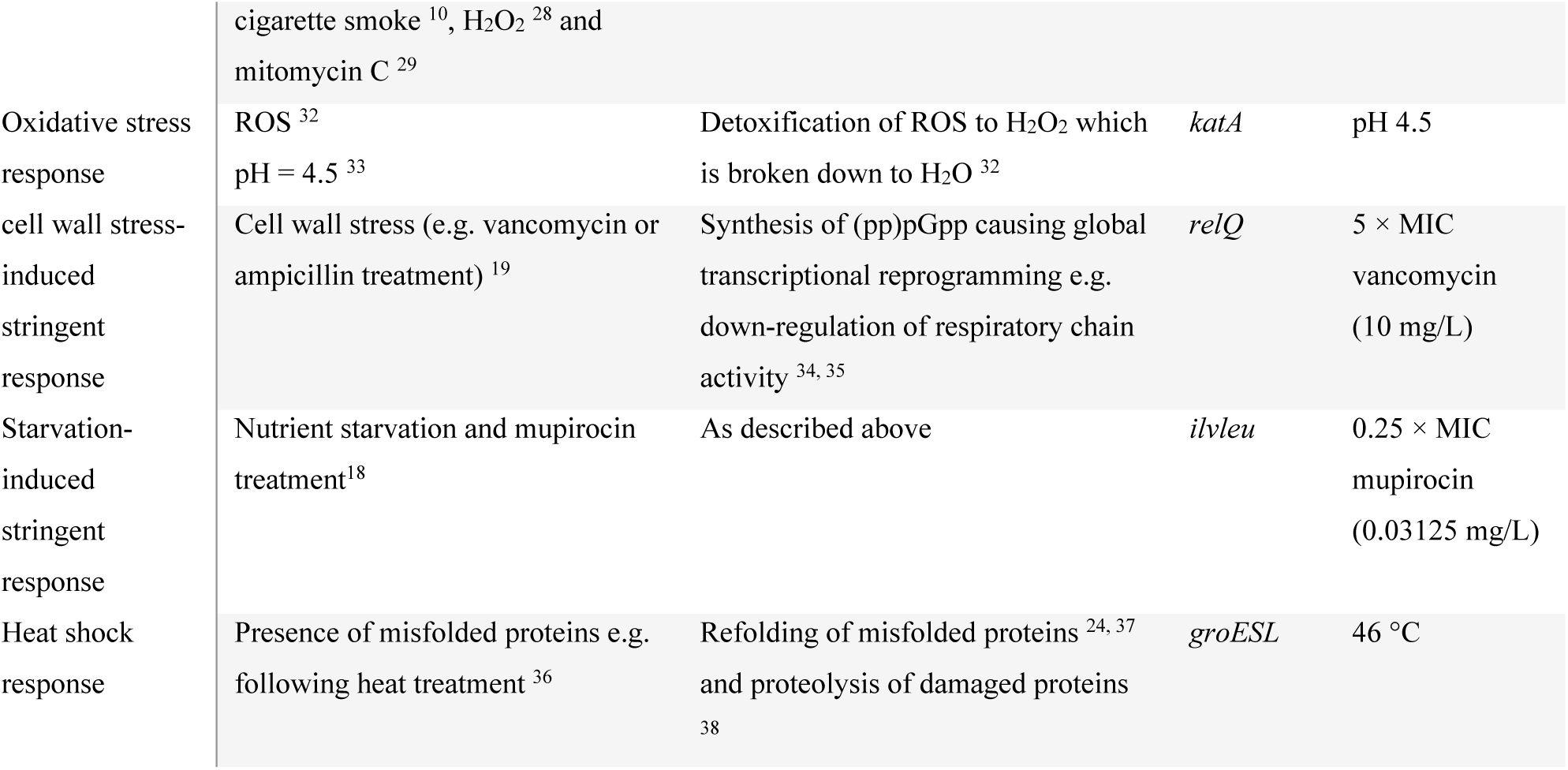
Details about stress responses investigated in this study.

Using these strains, we asked whether stress responses are activated by biofilm growth, by the host environment, or by both. We also investigated if any such activation is uniform across the population, and whether it coincides with a reduced rate of antibiotic killing. We find that biofilms grown in laboratory media show essentially no stress response activation, whereas biofilms of the same strain grown in human serum activate all five responses uniformly across the population. The effect also occurs in heat-inactivated serum, and a serum exposure as short as one hour is sufficient to abolish detectable killing by ciprofloxacin. The host environment, and not the biofilm growth mode, is thus the dominant driver of stress response activation in this system — with direct implications for the predictive value of laboratory biofilm models.

## Materials and methods

### Bacterial strains and growth conditions

*S. aureus* ATCC 29213 was cultured in tryptic soy broth (TSB, Sigma Aldrich, T8907) at 37 °C, 180 rpm. Reporter strains were cultured in TSB with 10 mg/L trimethoprim (Sigma Aldrich, T7883) overnight and then grown to exponential phase without trimethoprim prior to each experiment. Biofilms were grown in tryptic soy broth (Sigma Aldrich, 22092).

### Generation of reporter strains

To examine the expression of stress responses, five reporter systems were constructed by fusing the promoter (400 bp upstream of +1) of the genes, *recA, katA, relQ, ilvleu* or *groESL*, with *gfp_uvr_*and inserting the sequence in the shuttle vector pKK30. See overview of strains in Table 1. The sequence of each gene fusion is shown in Table S1, and all reporter strains have been deposited and are available through Addgene (deposit # 85764).

For the generation of reporter strains, *Escherichia coli* JM109λpir was used for plasmid storage and replication and grown in LB broth (Miller, 1102855000, Sigma Aldrich). Furthermore, *S. aureus* RN4220 was used to methylate plasmid DNA to prevent digestion by the restriction modification system in *S. aureus* ATCC 29213.

The fused promoter-*gfp_uvr_* was ordered from GenScript and delivered in pUC57. The promoter-*gfp_uvr_* construct was amplified from pUC57 using Phusion Green Hot Start II High-Fidelity Master Mix (ThermoFisher, F566S) and cycled in a BioRad T 100 Thermal cycler with settings: 98 °C for 30 s, 30 × (98 °C for 10 s, 59.6 °C for 30 s, 72 °C for 15 s), 72 °C for 10 min. Primers were designed with overhangs for Gibson cloning and lower case letters denote the overhangs. Forward primers: 5’-gctagcctaggagctCGATGTTCTAAAGGGTATGATTAATC-3’(*recA*), 5’-cggccgcta-gcctaggagctTAACAAGATAAGCGAGTATAGCG-3’ (*katA*), 5’-gctagcctaggagctGAGCTCGAACTTCGTACAATTC-3’ (*relQ*), 5’-cggccgctagcctaggagctCCTATATTATGCTTTTCATTCATAAAAATG-3’ (*ilvleu*), 5’-cggccgctagcctag-gagctGTGCTAAACTTTAGGTTTTTTAAGG-3’ (*groESL*). Reverse primer: 5’-ggatccccgggtaccgagctTTATTTATAT-AATTCATCCATACCATGTG-3’. Primers were designed using NEBuilder Assembly Tool (v 2.7.1, New England Biolabs). The constructs were inserted into pKK30 using Gibson cloning following the manufacturers protocol (Gibson Assembly Master Mix, New England Biolabs, E2611L). The pKK30-promoter construct was transformed into chemically competent *E. coli* JM109λpir through heat shock transformation for storage and replication, extracted using GenElute plasmid miniprep kit (Sigma Aldrich, PLN70) and transformed into electrocompetent *S. aureus* RN4220 as described elsewhere ^39^ using electroporation at 2.5 kV, 25 µF, 200 Ω, 4.5 ms. Finally, the pKK30-promoter construct was extracted using GenElute plasmid miniprep kit and transformed into electrocompetent *S. aureus* ATCC 29213 using electroporation at 2.5 kV, 25 µF, 200 Ω, 4.5 ms with subsequent growth at 37 °C for 1 h and plating on Tryptic Soy Agar with 10 µg/mL trimethoprim (Sigma Aldrich, T7883). Positive transformants were confirmed using colony PCR and sequenced to verify a correct construct.

### Minimum inhibitory concentration determination

The MIC of ciprofloxacin and vancomycin was determined using broth dilution in TSB. Briefly, an overnight culture of *S. aureus* ATCC 29213 was added to 2-fold serially diluted antibiotics in TSB in 96-well plates to obtain a starting concentration of 5 × 10^5^ CFU/mL. 96-well plates were incubated at 50 rpm, 37°C overnight. Afterwards, MIC was determined as the lowest antibiotic concentration where growth was inhibited by analysing 96-well plates in a plate reader at 600 nm.

### Validation of reporter strains using microscopy

Reporter strains were grown overnight and subsequently diluted 1000-fold in order to grow to the mid-exponential phase (OD_600_ = 0.3-0.5). Stress responses were subsequently induced by 1 h treatment with: 2 × MIC ciprofloxacin (2 mg/L) for the SOS response (*recA*), 5 × MIC vancomycin (10 mg/L) for the cell wall stress-induced stringent response (*relQ*), 0.25 × MIC mupirocin (0.03125 mg/L) for the starvation-induced stringent response (*ilvleu*), pH 4.5 (TSB adjusted with HCl) for the oxidative stress response (*katA*), or up to 3 h incubation at 46 °C for the heat shock response (*groESL*). To prepare a stationary culture, the overnight cultures were diluted 10-fold in Phosphate Buffered Saline (PBS, PIER28354, Thermo Fisher). Stationary phase, exponential phase and induced samples were stained with 20 µM SYTO 60 (S11342, invitrogen) and imaged in a Zeiss LSM700 confocal laser scanning microscope (CLSM). A 488 nm laser was used for excitation of GFP and a 639 nm laser was used for excitation of SYTO 60.

### Cross-activation of stress responses

Reporter strains were grown overnight and subsequently diluted 1:1000 in order to grow to mid-exponential phase (OD_600_ = 0.3-0.5). Every reporter strain received treatment with 2 × MIC ciprofloxacin (2 mg/L), 5 × MIC vancomycin (10 mg/L), 0.25 × MIC mupirocin (0.03125 mg/L), pH 4.5 (TSB adjusted with HCl) and 3 h incubation at 46 °C. As a negative control, reporter strains were kept in the mid-exponential phase and remained untreated. The samples were transferred to a 96-well plate and bulk fluorescence was measured in a fluorescence microplate reader (CLARIOstar, BGM LABTECH) with GFP excitation at 488 nm. The measured relative fluorescence from the reporter strains receiving treatment was normalised to the untreated negative controls.

### Antimicrobial time-kill assays

An overnight culture of *S. aureus* 29213 WT was diluted 1:1000 in TSB and grown to mid-exponential phase (OD_600_ = 0.3-0.5. Samples were washed by centrifugation for 5 min at 13150 × *g*, the supernatant was removed and the pellet was resuspended in either TSB (untreated), TSB with 2 × MIC ciprofloxacin, TSB adjusted to pH = 4.5, TSB with 5 × MIC vancomycin, TSB with 0.25 × MIC mupirocin, TSB pre-warmed to 46°C or 100% serum and incubated for 1 h at 37 °C (or 46 °C for heat-treated samples). Subsequently, samples were washed by centrifugation at 13150 × *g*, the supernatant was removed and the pellets were resuspended in TSB (untreated), TSB with 5 × MIC ciprofloxacin, 5 × MIC vancomycin, 5 × MIC daptomycin, 5 × MIC dicloxacillin or 5 × MIC rifampicin for 2 h. CFU was enumerated to quantify survival before pre-treatment (T = 0 h), after pre-treatment (T = 1 h), and after treatment (T = 3 h).

### Stress response activation in human serum

Serum was prepared from blood (irreversibly anonymized) collected in BD Vacutainer tubes (VWRTM, 367896) by Aarhus University Hospital (AUH) blood bank, and prepared by centrifugation at 2000 × *g* for 15 min at 4°C. Serum from at least 6 patients were pooled to minimize sample variation and then aliquoted at 2-8°C, and stored at -80°C. Inactivated serum was also used and was prepared by heating it to 56°C for 30 min to inactivate the complement system.

Reporter strain were grown overnight, diluted 1:1000 in fresh TSB and allowed to grow to mid-exponential phase (OD_600_ = 0.3-0.5) for approx. 2 h with 0.1 mM HADA, a blue fluorescent *D*-amino acid that is incorporated into the peptidoglycan cell wall during bacterial growth (Bio-techne, 6647). Subsequently, samples were centrifuged for 5 min at 13150 × *g*, the supernatant was removed and pellets were resuspended in either TSB (untreated, 0% serum), 10% human serum in TSB or in 100% human serum and incubated for 1 h at 37 °C and 100 rpm orbital shaking. Afterwards, the reporter strains were washed by centrifugation for 5 min at 13150 × *g*, the supernatant was removed and the pellets were resuspended in PBS. Samples were mounted to glass slides, allowed to dry out, followed by adding a drop of antifade (Thermo Scientific, P36980) and putting on a coverslip. The samples were visualised by CLSM using a 405 nm laser to excite HADA and a 488 nm laser to excite GFP.

GFP fluorescence was quantified in Image J by first using the GFP signal to create a mask, and then quantify the average pixel intensity of the area inside the mask. The HADA image was used to assess if the mask correctly identified the area of all bacteria in the sample.

### Biofilm formation

Reporter strains were grown overnight, diluted 1:1000 in fresh TSB and allowed to grow to mid-exponential phase (OD_600_ 0.3-0.5). An 8-well Ibidi (Ibidi, 80826) µ-slide was pre-conditioned by adding 100% human plasma for 30 min before removal. The optical density of the exponential cultures was adjusted to OD_600_ = 0.1 in either TSB, TSB with 10% human serum or 100% human serum and the bacteria were added to the pre-conditioned wells and incubated for 30 min at 37 °C without shaking. Subsequently, the liquid was removed from the wells and fresh TSB, 10% serum in TSB or 100% serum was added and wells were supplemented with 0.1 mM HADA. Biofilms were incubated for up to 72 h with a change in media every 24 h as described above. Immediately before imaging, the liquid from each well was removed and replaced by PBS. The same CLSM settings were used as described in the section above.

### Neutrophil phagocytosis of reporter strains

For human neutrophil isolation, blood was collected in heparin-coated tubes (BD Vacutainer 367526) from healthy donors from Aarhus University Hospital, Denmark. Immediately after collection, 5 mL blood was transferred to 5 mL polymorph-prep density gradient medium (Alere, 1114683) and centrifuged at 500 × *g* for 35 min at 20 °C. Deceleration of the centrifuge rotation was performed without braking. Plasma and monocytes were removed and neutrophils were collected and mixed with Hanks balanced salt solution without Mg^2+^ and Ca^2+^ (HBSS-, Gibco, 14175095) and centrifuged at 400 × *g* for 10 min. The pelleted neutrophils were resuspended in ACK lysing buffer (Thermo Fisher Scientific, A1049201) for 3 min at room temperature in order to lyse any remaining erythrocytes. Subsequently, neutrophils were washed by centrifugation for 5 min at 300 × *g* at 20 °C, removing the supernatant and resuspending in HBSS-before centrifuging for 10 min at 120 × *g,* removing supernatant and resuspending in HBSS-. Neutrophils were counted and the concentration of neutrophils was adjusted to 5 × 10^6^ PMNs/mL in HBSS+ (HBSS with Mg^2+^ and Ca^2+^, Gibco, 24020117).

Overnight cultures of reporter strains had been prepared the day before with 0.1 mM HADA. The grown cultures were adjusted to OD_600_ = 0.1 when mixed with the neutrophils and phagocytosis was initiated by adding 10% human serum for 2 h at 37°C and 5% CO_2_. Subsequently, 2 mM EDTA was added to prevent clumping of neutrophils and samples were fixated in 4% paraformaldehyde for 45 min before being resuspended in PBS with 3% bovine serum albumin (BSA, Sigma Aldrich, A3294). Neutrophils were stained in a 1:50 ration of recombinant Alexa Fluor 594 anti-CD16 antibody (Abcam, ab302799) and samples were imaged using CLSM. CLSM settings: GFP (ex. 488 nm, em 490-580 nm), HADA (ex. 405 nm, em. 405-480 nm), anti-CD16 (ex. 555 nm, em. 580-750 nm) with a beam splitter placed at 480 nm to prevent cross-talk. Non-phagocytosed/free reporter strains (negative control) received the same treatment as the phagocytosed *S. aureus* but without exposure to neutrophils. GFP signal intensity was quantified as described above.

### Data analysis and statistics

Data was tested using a t-test for single comparisons and one-way ANOVA for multiple comparisons with post-hoc uncorrected Fisher’s LSD. In text, data is displayed as mean ± standard deviation. GraphPad Prism was used for all statistical analyses (v. 9.5.1 (733) for Windows, GraphPad Software, San Diego, California USA, www.graphpad.com).

## Results

### Fluorescent transcriptional reporters resolve activation of five stress responses in single cells

To follow stress response activation in individual *Staphylococcus aureus* cells, we constructed five transcriptional reporters by fusing the promoter region of a representative gene to GFPuvr, a stable and strongly fluorescent variant of GFP ^23^, and inserting each fusion into the shuttle vector pKK30. The reporters monitor the SOS response (*recA*), the oxidative stress response (*katA*), the cell wall stress- and starvation-activated branches of the stringent response (*relQ* and *ilvleu*, respectively), and the heat shock response linked to misfolding of proteins (*groESL*). The stress reported by each construct, the rationale for the choice of gene, and the condition used to induce it are given in Table 1, and the minimum inhibitory concentrations of the antibiotics used throughout the study are listed in Table 2. All five reporter strains have been deposited at Addgene, and because the constructs are carried on a shuttle plasmid rather than integrated into the chromosome, they can be introduced into other strains, including clinical isolates.

**Table 2.** Minimum Inhibitory Concentrations of relevant antimicrobials.

| Antibiotic | MIC (mg/L) |
| --- | --- |
| Ciprofloxacin | 1 |
| Vancomycin | 2 |
| Mupirocin | 0.125 |
| Dicloxacillin | 0.125 |
| Rifampicin | 0.008 |
| Daptomycin | 2 |

Throughout this study, bacteria were counterstained with the fluorescent D-amino acid HADA, which is incorporated into peptidoglycan during growth and allows cells to be located independently of GFP signal. HADA did not itself activate any of the five reporters (Fig. S1), and an isogenic strain carrying the empty pKK30 plasmid was imaged alongside the reporter strains in every experiment to define the level of background fluorescence.

We first confirmed that each reporter responded to its cognate stress. Exponentially growing cultures were exposed for 1 h to 2 × MIC ciprofloxacin, TSB adjusted to pH 4.5, 5 × MIC vancomycin, 0.25 × MIC mupirocin, or a temperature of 46 °C, and imaged by confocal microscopy alongside untreated exponential a phase cultures (Fig. 1). Each condition activated its intended reporter. In stationary phase, the SOS, oxidative stress, starvation-induced stringent and heat shock reporters were all activated (Fig. S2), consistent with the multiple stresses experienced by non-growing cultures in spent medium. Exponential phase cultures were therefore used for all subsequent experiments.

**Fig. 1.**
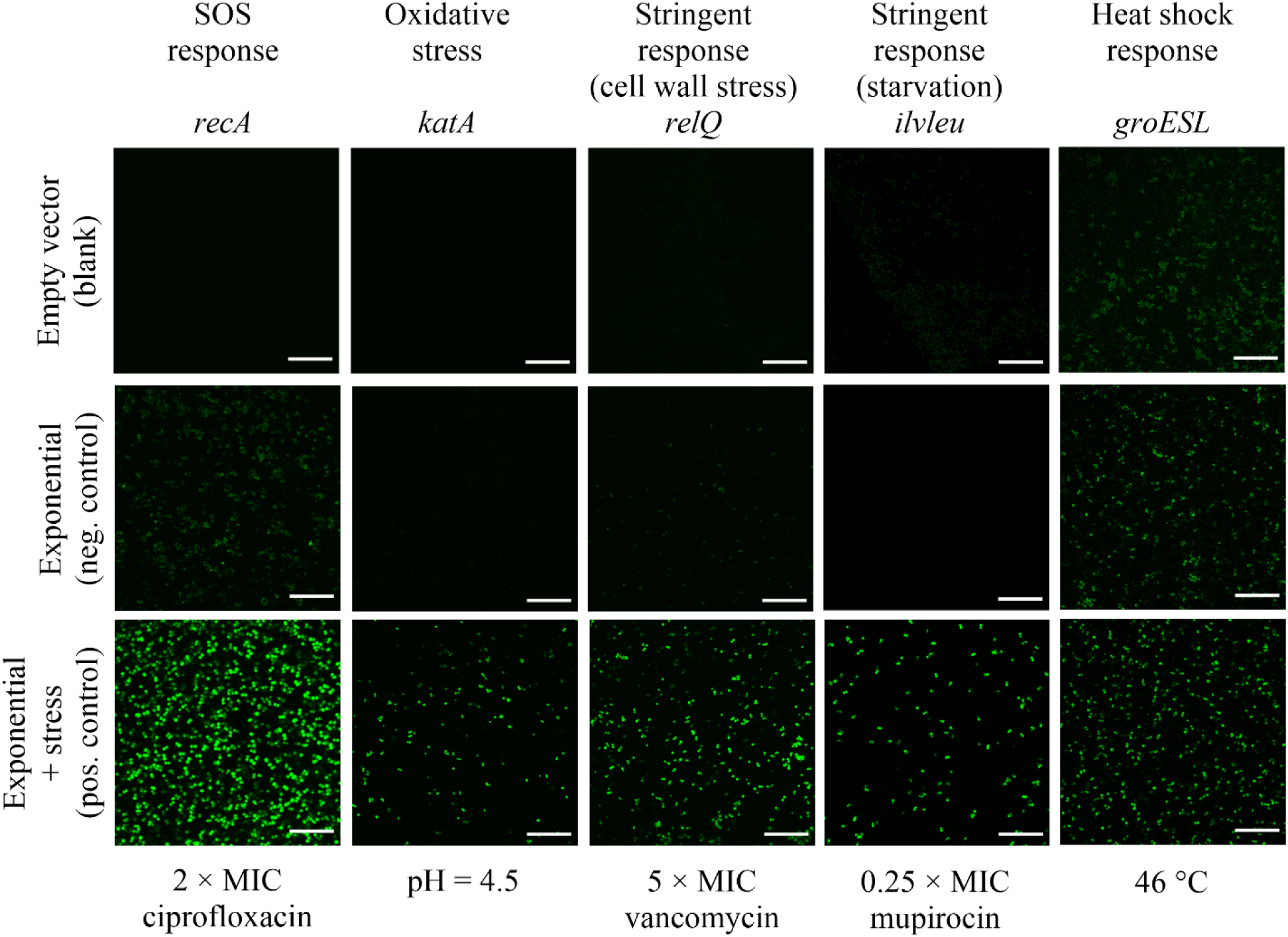
Activation of stress responses in exponentially growing cultures. *S. aureus* 29213 pKK30 P*recA-gfp* (SOS response), *S. aureus* 29213 pKK30 P*katA-gfp* (oxidative stress response), *S. aureus* 29213 pKK30 P*relQ-gfp* (stringent response/cell wall stress), *S. aureus* 29213 pKK30 P*ilvleu-gfp* (stringent response/starvation), *S. aureus* 29213 pKK30 P*groESL-gfp* (heat shock response) and *S. aureus* 29213 pKK30 (empty plasmid control) were grown overnight and subsequently diluted 1000-fold in TSB + 0.1 mM HADA in order to grow to mid-exponential phase. Each stress response was specifically induced by stressing the exponential cultures for 1 h with 2 × MIC ciprofloxacin (SOS response), TSB adjusted to pH = 4.5 with HCl (oxidative stress response), 5 × MIC vancomycin (stringent response/cell wall stress), 0.25 × MIC mupirocin (stringent response/starvation) and 46 °C (heat shock response). Subsequently cultures were visualised by CLSM using 405 nm laser to excite HADA (data not shown) and a 488 nm laser to excite GFP. Scale bar: 20 µm. n = 3 biological replicates.

In exponentially growing cultures, four of the five reporters were at or close to the fluorescence level of the empty-plasmid control. The SOS reporter was the exception, showing a low but detectable signal in exponential phase (Fig. 1). We therefore used exponentially growing cultures as the reference condition throughout, while noting that a basal level of recA expression is present in this state and that activation of the SOS reporter should be read against that baseline rather than against zero.

An observation common to all five reporters was that activation was not restricted to a subset of cells. Where a stress condition activated a reporter, GFP fluorescence was apparent in essentially all cells present in the field of view, rather than in a scattered minority against an unlabelled background (Fig. 1). This distinction is central to the interpretation of the experiments that follow, because a stress response mounted by an entire population and one mounted by a small subpopulation have different consequences for how a population is killed by antibiotics.

Because environmental conditions rarely impose a single stress, we next investigated the specificity of each induction condition. Each of the five reporter strains was exposed to each of the five conditions, and bulk fluorescence was measured in a plate reader and normalised to the untreated control of the same strain (Fig. 2). Ciprofloxacin, vancomycin, mupirocin and 46 °C each activated only the reporter they were intended to activate. Exposure to pH 4.5 was different: in addition to the oxidative stress reporter, it significantly activated both the SOS reporter and the cell wall stress-induced stringent reporter (Fig. 2B).

**Fig. 2.**
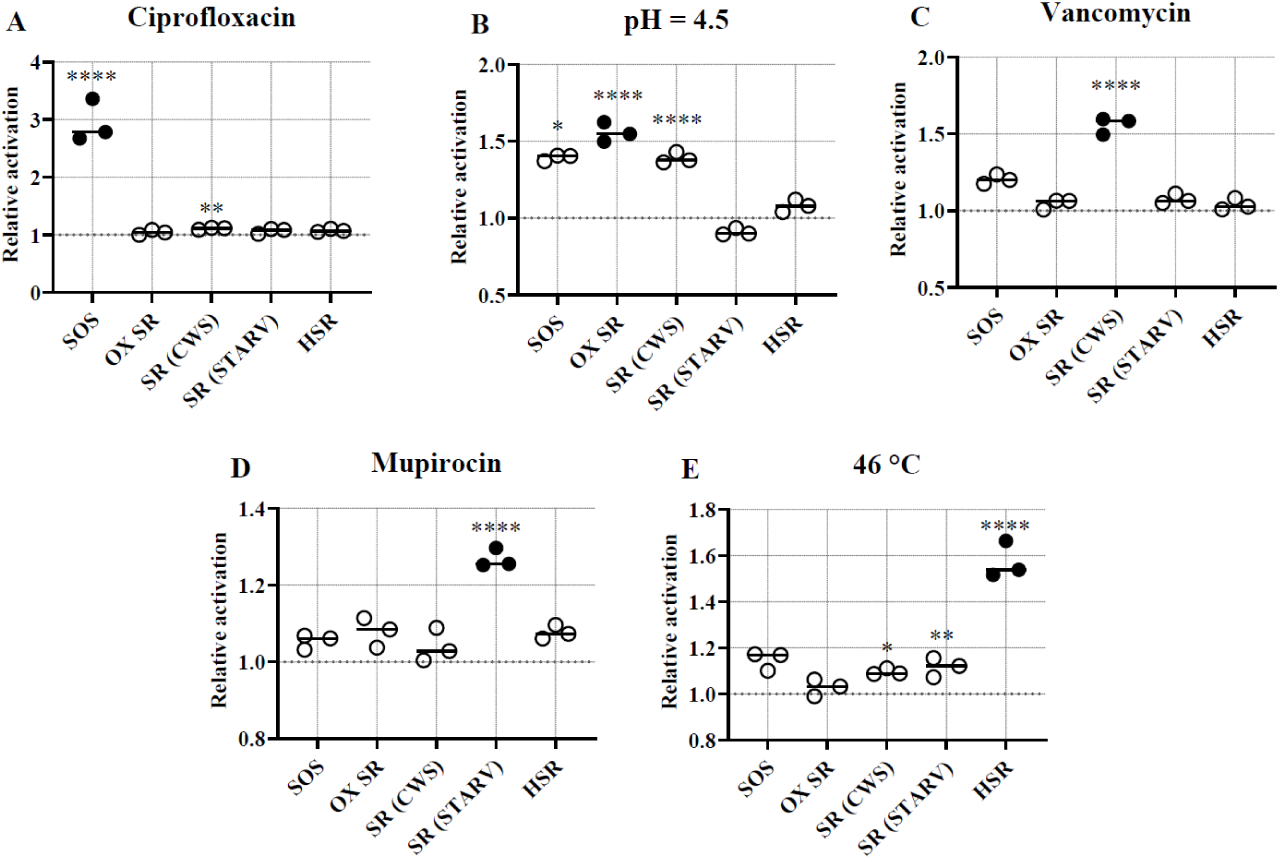
Cross activation of stress responses during specific induction. Stress response reporter strains and the empty plasmid control strain were grown to mid-exponential phase and subjected to the array of specific stressors used to induce each stress response. Each reporter strain was either left untreated or treated with 2 × MIC ciprofloxacin (A), TSB adjusted to pH = 4.5 with HCl (B), 5 × MIC vancomycin (C), 0.25 × MIC mupirocin (D) and 46 °C (E). Fluorescence was measured at 488 nm excitation and 510 nm emission. Relative fluorescence units for each reporter strain was normalised to the mean fluorescence of the untreated control of that reporter strain (corresponding with relative activation = 1). Black circles denote the stress responses which are intended to be activated from the given treatment. For statistical testing: The fluorescence from each treated and untreated sample was normalised to the fluorescence from the empty plasmid control. Treated samples were then compared to untreated samples using ordinary one-way ANOVA with Dunnett’s multiple comparisons test. **P = 0.0021, ****P < 0.0001. n = 3 biological replicates.

We therefore refer to the pH 4.5 condition as acid stress rather than as an oxidative stress condition for the remainder of this study. Because acid exposure activates three of the five responses, any effect observed after acid treatment cannot be attributed to the oxidative stress response specifically.

### Biofilm formation in laboratory media does not activate stress responses

The tolerance of biofilm infections to antibiotics is conventionally attributed to the biofilm mode of growth itself. Mass transfer limitation within the biofilm is presumed to generate gradients of oxygen and nutrients, and the starvation and growth arrest that follow are presumed to render cells in the deeper layers refractory to killing. If this model is correct, biofilm growth should be sufficient to activate stress responses, and particularly the starvation-activated branch of the stringent response.

We tested this directly. Reporter strains were grown for 24 h as biofilms in TSB in plasma-conditioned wells and imaged by confocal microscopy (Fig. 3). Dense microcolonies formed under these conditions, but none of the five reporters showed appreciable activation that could be distinguished from the empty-plasmid control strain imaged with identical settings (Fig. S3). Biofilm growth in rich laboratory medium is therefore not by itself sufficient to activate any of the five stress responses examined here.

**Fig. 3.**
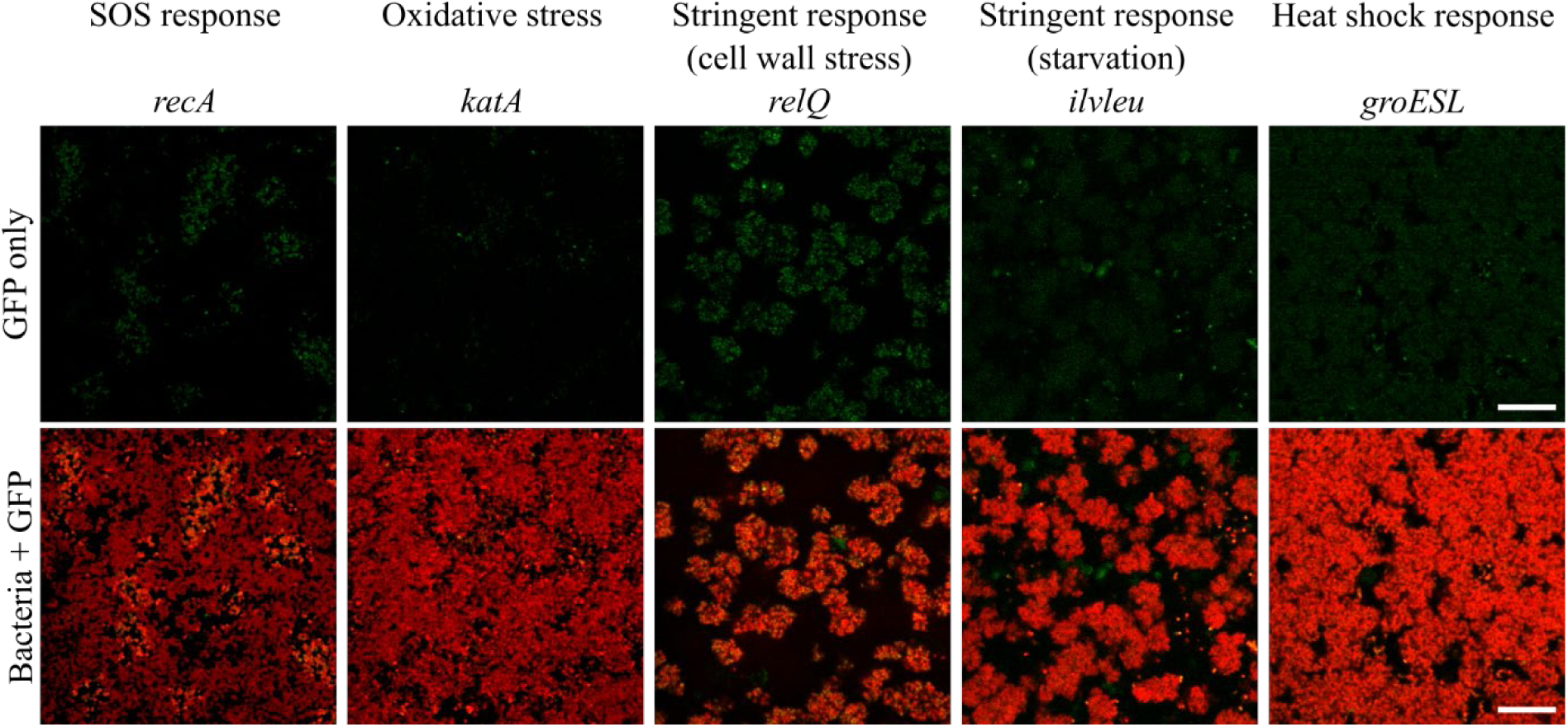
Biofilms grown in laboratory media do not activate stress responses. Reporter strains carrying *recA–gfpuvr* (SOS response), *katA–gfpuvr* (oxidative stress response), *relQ–gfpuvr* (stringent response, cell wall stress), *ilvleu–gfpuvr* (stringent response, starvation) or *groESL–gfpuvr* (heat shock response) were grown to mid-exponential phase, resuspended in TSB containing 0.1 mM HADA, and inoculated into wells pre-conditioned with 100% human plasma. After 24 h of growth, biofilms were washed with PBS and imaged by confocal laser scanning microscopy. Green, GFP; red, HADA (peptidoglycan cell wall). Images are 2D sections acquired from approximately the middle of the biofilm. n = 3 biological replicates. Scale bar: 20 µm. The scale is identical in all images

### The host environment activates all five stress responses across the biofilm population

We next grew biofilms of the same reporter strains under host-relevant conditions, replacing the laboratory medium with 100% human serum while keeping all other aspects of the assay unchanged. In contrast to biofilms grown in TSB, biofilms grown in serum activated all five reporters (Fig. 4). Activation of the SOS response, the oxidative stress response, the cell wall stress-induced stringent response, the starvation-induced stringent response and the heat shock response was pronounced. It should be noted that acquisition settings had to be optimised for each strain, so GFP fluorescence is comparable within strains across the different growth media (TSB vs serum), but it is not directly comparable between strains.

**Fig. 4.**
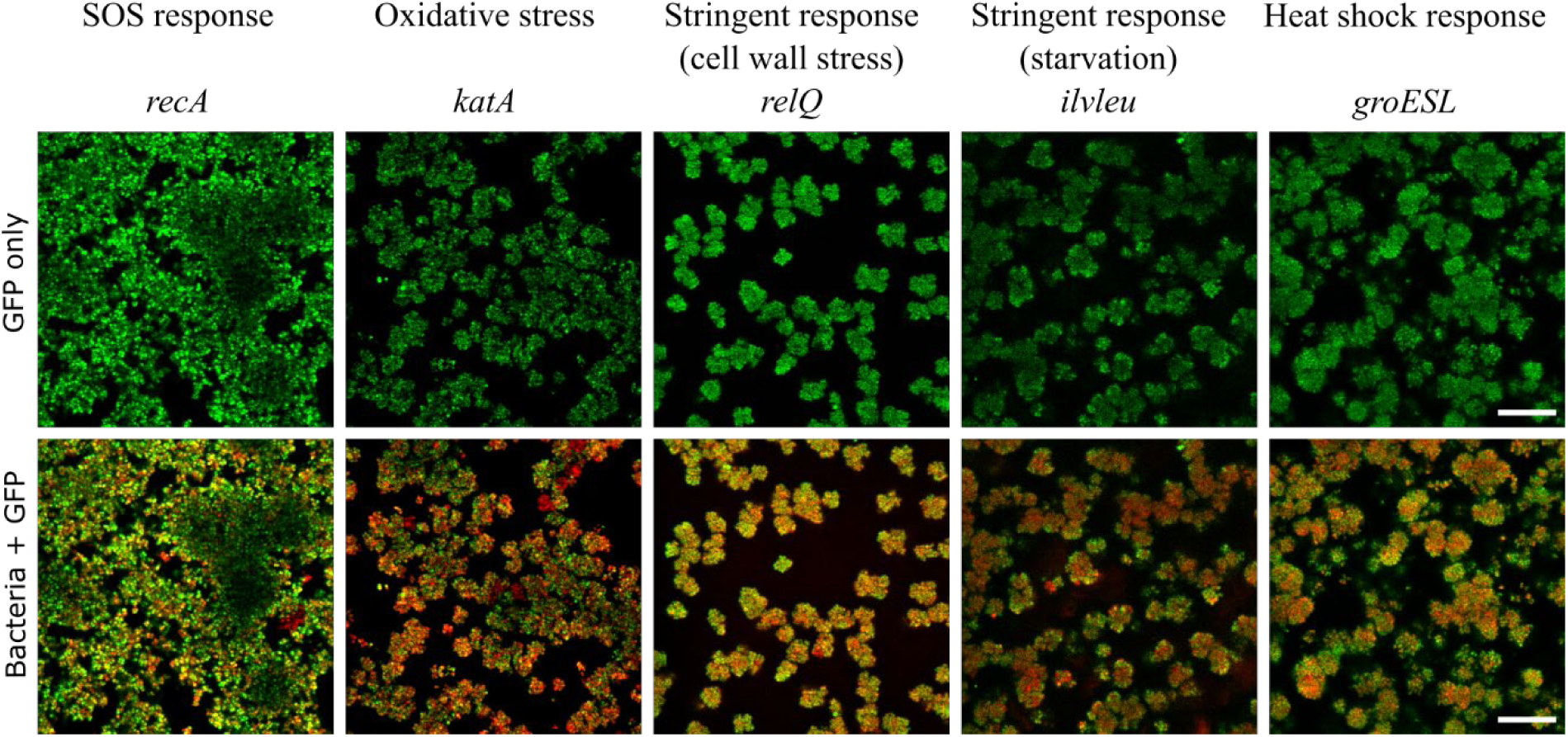
Biofilms grown in human serum activate all five stress responses across the population. Reporter strains were grown to mid-exponential phase, resuspended in 100% human serum containing 0.1 mM HADA, and inoculated into wells pre-conditioned with 100% human plasma. After 24 h of growth, biofilms were washed with PBS and imaged by confocal laser scanning microscopy. Green, GFP; red, HADA (peptidoglycan cell wall). Images are 2D sections acquired from approximately the middle of the biofilm. Acquisition settings were identical to those used in Fig. 3. The corresponding empty-plasmid control imaged with these settings is shown in Fig. S2. n = 3 biological replicates. Scale bar: 20 µm.

Activation was not confined to a subpopulation or to a particular region of the microcolonies. GFP fluorescence was apparent throughout the biofilms, in essentially all cells identified by HADA counterstaining (Fig. 4), indicating that the serum environment activates these responses across the population rather than in a distinct subset of cells.

We noted that the fluorescence intensity of both HADA and GFP was reduced towards the centre of large microcolonies, e.g. visible in image showing the *recA* reporter (Fig. 4). This may reflect light scattering within the biofilm, or reduced GFP maturation and HADA incorporation under the locally anoxic conditions expected in the interior. Because the effect was seen for all five reporter strains and affected the HADA counterstain as well as the GFP signal, we do not interpret it as reduced stress response activation in the biofilm interior.

The effect of serum on stress response activation was concentration-dependent. Biofilms grown in 10% human serum diluted in TSB showed no visible activation of any reporter (Fig. S4), resembling biofilms grown in TSB alone rather than those grown in undiluted serum. This has a practical consequence that we exploit below: 10% serum can be used to opsonise bacteria for phagocytosis without itself activating the reporters.

To ask whether the stress-inducing activity of serum depended on heat-labile components, we grew biofilms in serum that had been heat-inactivated at 56 °C for 30 min, a treatment that inactivates complement. Heat-inactivated serum activated the reporters in the same manner as native serum (Fig. S5). The factors responsible are therefore not complement proteins or other heat-labile serum enzymes, but are more likely to reside in the nutritional composition of serum, including its restriction of free iron and zinc, or in heat-stable antimicrobial components.

Taken together, these results separate two variables that are usually confounded. Biofilms grown in laboratory medium or serum share the same mode of growth and incubation period, differing only in the medium. It is therefore the host environment, and not the biofilm mode of growth, that drives stress response activation in this system.

### Brief exposure to human serum activates the stringent response and abolishes killing by ciprofloxacin

The strong activation of stress responses in serum raises the question of whether the host environment also renders *S. aureus* less susceptible to antibiotics. We addressed this by exposing exponentially growing wild-type bacteria to 100% human serum for 1 h, and then challenging them for 2 h with ciprofloxacin, vancomycin, daptomycin, dicloxacillin or rifampicin, each at 5 × MIC, in TSB (Fig. 5A). Killing was quantified as the change in viable count during the antibiotic challenge and compared with cultures that had not been pre-exposed to serum (Fig. 5B-H). We used a 1 h exposure for consistency with the experiments described in the following section, recognising that this is considerably shorter than the 24 h of serum exposure used in the biofilm experiments.

**Fig. 5.**
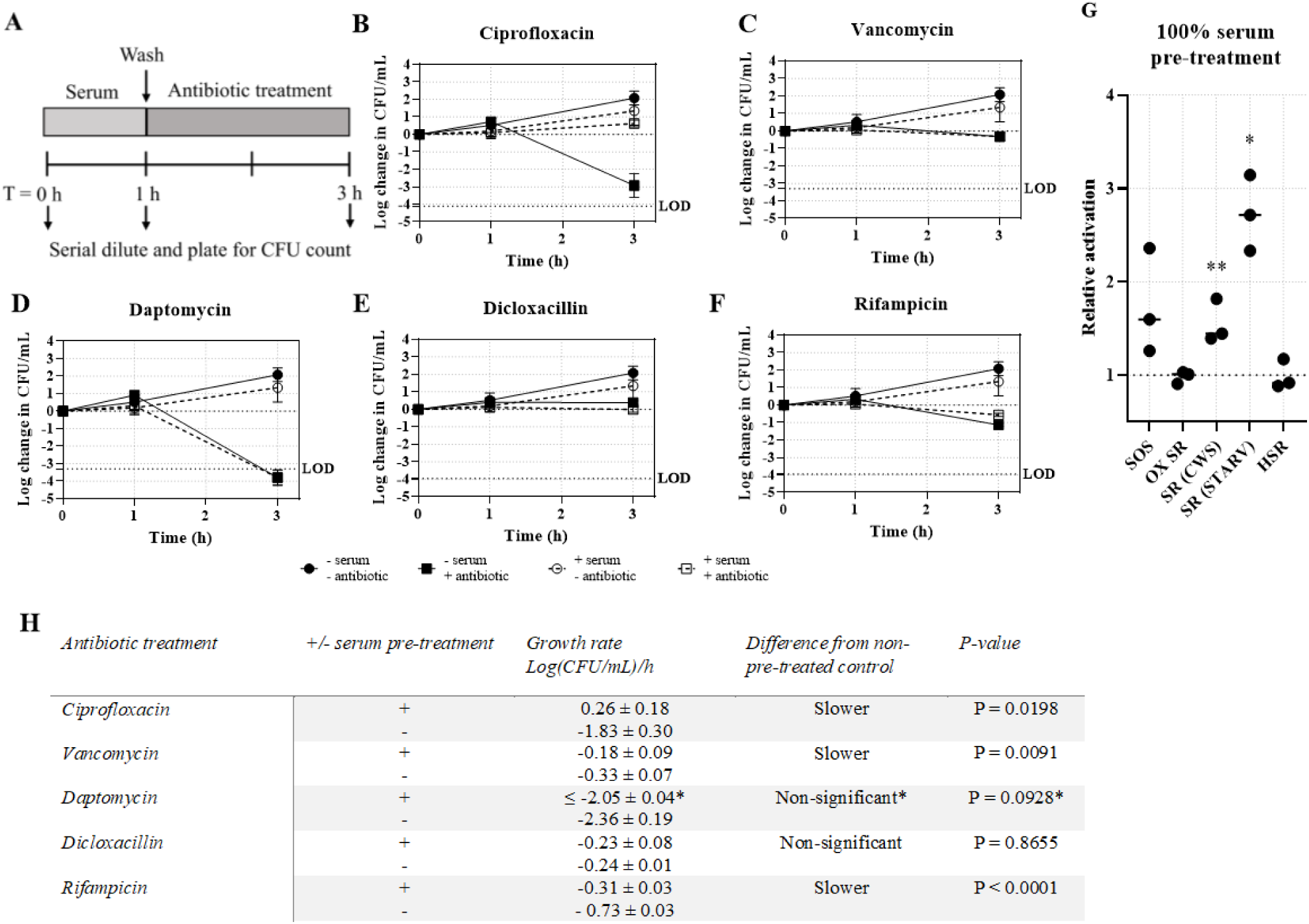
Antibiotic treatment following pre-treatment with human serum. *S. aureus* 29213 WT was grown to mid-exponential phase and subsequently received 1 h pre-treatment in 100% human serum followed by 2 h treatment selected antibiotics where CFU was enumerated a T = 0, 1 and 3 h (A). Antibiotic treatment was performed using 5 × MIC ciprofloxacin (B), 5 × MIC vancomycin (C), 5 × MIC daptomycin (D), 5 × MIC dicloxacillin (E) or 5 × MIC rifampicin (F). Samples without serum pre-treatment (“-serum”) and samples without ciprofloxacin treatment (“-antibiotic”) were incubated in TSB at 37°C. n = 3 biological replicates for samples with antibiotic treatment and n = 9 biological replicates for samples without antibiotic treatment. Means are shown as symbols and standard deviations are shown as error bars. Where no error bars are present, the standard deviation is too small to be seen. LOD: limit of detection. (G) Stress response reporter strains were grown to mid-exponential phase with 0.1 mM HADA for counterstaining and treated for 1 h in 100% serum or left untreated in TSB followed by washing and resuspending in PBS. The reporter strains were mounted on a glass slide with antifade and visualised by CLSM using a 405 nm laser to excite HADA (to locate cells) and a 488 nm laser to excite GFP. GFP fluorescence intensities for the serum-treated samples were normalised to the GFP intensities for the untreated samples to achieve the relative activation of stress responses in serum. n = 3 biological replicates. Statistical testing was performed using a one-way ANOVA. *P = 0.0102, **P = 0.0064. (H) Growth rates (negative values indicate killing) in response to antibiotic treatment, calculated for pre-treated and untreated samples from CFU counts at T1 and T3. **\***The CFU count after daptomycin treatment was below the limit of detection and the death rate cannot be determined precisely. Statistical testing was performed for the displayed death rate. Statistical analysis: unpaired t-test. n = 3 biological replicates biological replicates for the untreated samples. Shown are the mean death rates ± standard deviations.

Serum exposure alone had no antimicrobial effect. Its effect on subsequent antibiotic killing was, however, striking and largely specific to ciprofloxacin. Bacteria that had not been exposed to serum were killed rapidly by ciprofloxacin, at a rate of 1.83 ± 0.30 log (CFU/mL)/h, whereas bacteria pre-exposed to serum showed no detectable killing at all; viable counts increased slightly over the challenge period (Fig. 5B, 5H). A 1 h exposure to serum was therefore sufficient to abolish killing by ciprofloxacin entirely.

The effect on the other antibiotics was small or absent. Serum pre-exposure produced a statistically significant but minor reduction in the rate of killing by vancomycin and by rifampicin (Fig. 5C, 5F, 5H), in both cases a change of less than half a log per hour, which we consider unlikely to be of clinical consequence. Killing by daptomycin was not significantly affected (Fig. 5D), but it should be noted that the CFU count at the end of the treatment was below the detection limit, making the quantification inaccurate. Killing by dicloxacillin was likewise unaffected (Fig. 5E), but this comparison is uninformative, because dicloxacillin at 5 × MIC produced almost no killing even in the absence of serum pre-treatment (Fig. 5H).

To determine which responses were activated over this short exposure, reporter strains were treated in parallel under identical conditions and imaged by confocal microscopy (Fig. 5G). One hour in serum significantly activated both branches of the stringent response, i.e. the cell wall stress-activated branch (*relQ*) (P = 0.0102) and the starvation-activated branch (*ilvleu*) (P = 0.0064). Activation of the remaining reporters did not reach significance over this interval, which is considerably shorter than the 24 h of serum exposure that activated all five responses in biofilms (Fig. 4). A brief encounter with the host environment is thus sufficient to activate the stringent response and to abolish ciprofloxacin killing, and the broader activation seen in biofilms grown in serum for 24 h suggests that 1 h of exposure underestimates the effect of the host environment.

### Stress responses that arrest growth are associated with slower killing by ciprofloxacin

Having established that serum both activates stress responses and confers ciprofloxacin tolerance, we asked which individual responses could plausibly account for that tolerance. Individual responses were activated in the wild-type strain using the conditions validated in Fig. 1. The bacteria were challenged with their specific stress response activators for 1 h (the pre-treatment) before measuring the subsequent growth rate by CFU counts at the 1 h and 3 h mark during incubation with or without ciprofloxacin at 5 × MIC concentration (Fig. 6A). The death rate (i.e. negative growth rate) during ciprofloxacin exposure was compared for activated vs non-activated cultures, revealing if activation of a specific stress response provides protection against the antimicrobial activity of ciprofloxacin (Fig. 6G).

**Fig. 6.**
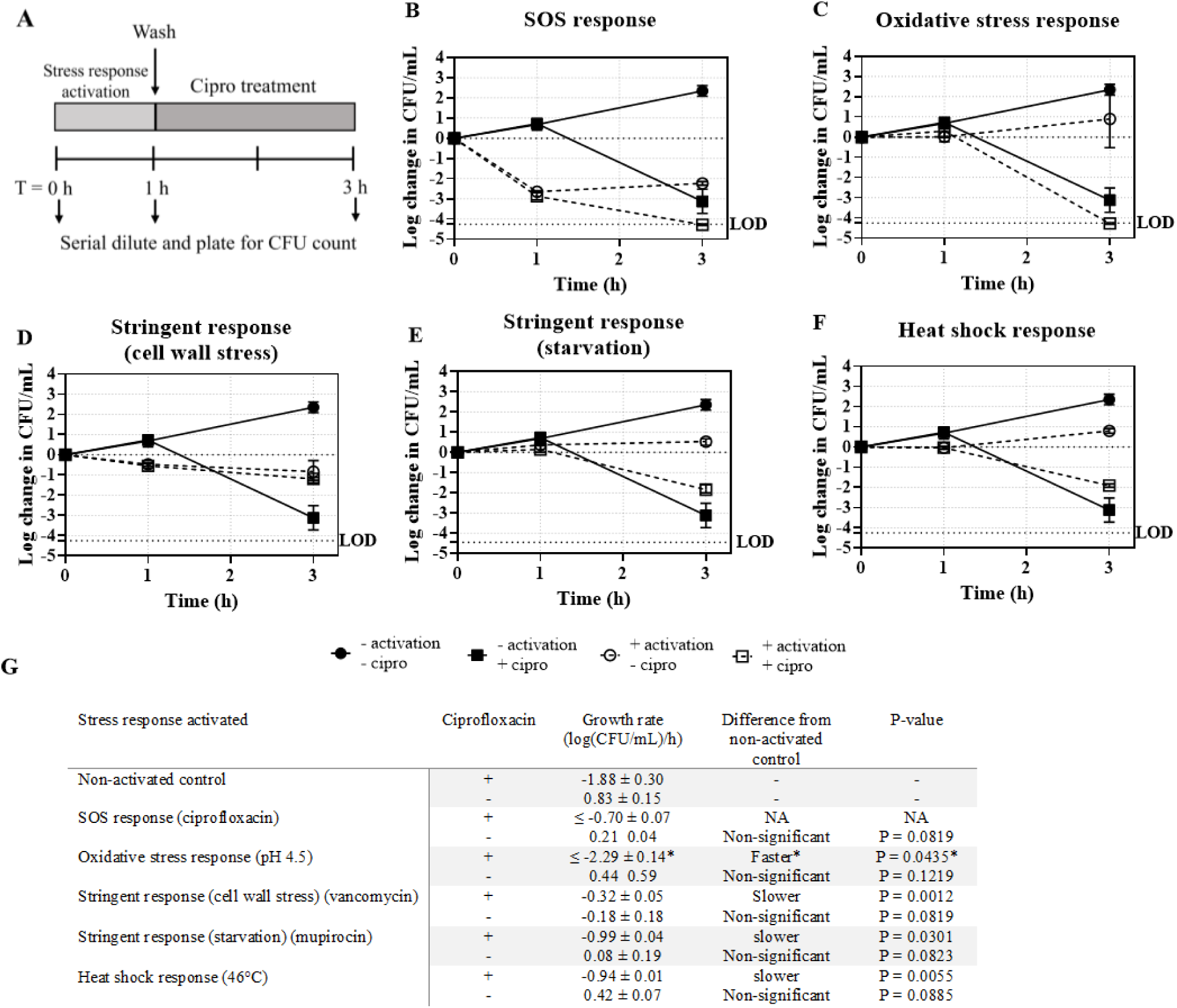
Antimicrobial effect of ciprofloxacin following stress response activation. *S. aureus* 29213 WT was grown to mid-exponential phase and subsequently received 1 h pre-treatment followed by 2 h treatment with 5 × MIC ciprofloxacin (+ cipro) where CFU was enumerated a T = 0, 1 and 3 h (A). Stress response activation through pre-treatments (+ activation) were performed with either 2 × MIC ciprofloxacin to activate the SOS response (B), TSB adjusted to pH = 4.5 to activate the oxidative stress response (C), 5 × MIC vancomycin to activate the stringent response related to cell wall stress (D), 0.25 × MIC mupirocin to activate the stringent response related to starvation (E) and 46 °C to activate the heat shock response (F). Samples without activation of stress responses (-activation) and samples without ciprofloxacin treatment (-cipro) were incubated in TSB at 37°C. n = 3 biological replicates for pre-treated samples and n = 12 biological replicates for samples without pre-treatment. Means are shown as symbols and standard deviations are shown as error bars. Where no error bars are present, the standard deviation is too small to be seen. LOD: limit of detection. (G) Death rates in response to antibiotic treatment, calculated for pre-treated and untreated samples from CFU counts at T1 and T3. *The survival reached below the limit of detection and the death rate cannot be determined precisely. Statistical testing was performed for the displayed death rate. Statistical analysis: unpaired t-test. n = 3 biological replicates biological replicates for the untreated samples. Shown are the mean death rates ± standard deviations. NA: The survival reached below the limit of detection and the death rate cannot be determined precisely. Statistical testing was not performed. *Statistical testing was performed for the death rate displayed. As the CFU concentration was below the limit of detection, the death rate may be even higher.-: The untreated control.

Activation of either branch of the stringent response reduced the rate of ciprofloxacin killing. Pre-treatment with vancomycin, which activates the cell wall stress-induced branch via *relQ*, reduced the death rate from 1.88 ± 0.30 to 0.32 ± 0.05 log (CFU/mL)/h (Fig. 6D, 6G), and pre-treatment with mupirocin, which activates the starvation-induced branch, reduced it to 0.99 ± 0.04 log (CFU/mL)/h (Fig. 6E, 6G). Pre-treatment at 46 °C, which activates the heat shock response, reduced the death rate to −0.94 ± 0.01 log (CFU/mL)/h (Fig. 6F, 6G).

Acid stress had the opposite effect, increasing the rate of killing to at least −2.29 ± 0.14 log (CFU/mL)/h (Fig. 6C, 6G). As established above, pH 4.5 activates three of the five responses, so this increase in susceptibility cannot be attributed to the oxidative stress response specifically. Pre-treatment with ciprofloxacin to activate the SOS response killed more than 99.9% of the population before the challenge began, leaving too few survivors for the subsequent killing to be quantified, and this condition was therefore uninformative (Fig. 6B, 6G).

One feature of these experiments constrains their interpretation. Each pre-treatment that reduced the rate of ciprofloxacin killing also arrested growth, and growth remained arrested for the duration of the antibiotic challenge (Fig. 6G, untreated rows). Activation of the stress response and cessation of growth are therefore completely confounded in this experiment, and the association we observe is between growth-arresting stress conditions and slower killing, rather than between a specific signalling pathway and slower killing. The same caveat applies to the serum experiments in the preceding section, where the stringent response was activated.

### Phagocytosis by neutrophils activates the oxidative stress and stringent responses

Serum is only one of the host environments *S. aureus* encounters. The bacterium is also taken up by phagocytes and can survive within them, where it is exposed to reactive oxygen species and low pH and is shielded from many antibiotics. We therefore asked whether the reporters were activated inside human neutrophils.

Neutrophils were isolated from whole blood and mixed with reporter strains, and phagocytosis was initiated by adding human serum to a working concentration of 10%, a concentration that does not itself activate the reporters (Fig. S3). After 2 h, samples were fixed and imaged by confocal microscopy, and GFP intensity in phagocytosed bacteria was compared with that of bacteria subjected to identical treatment, including the same serum exposure, in the absence of neutrophils (Fig. 7).

**Fig. 7.**
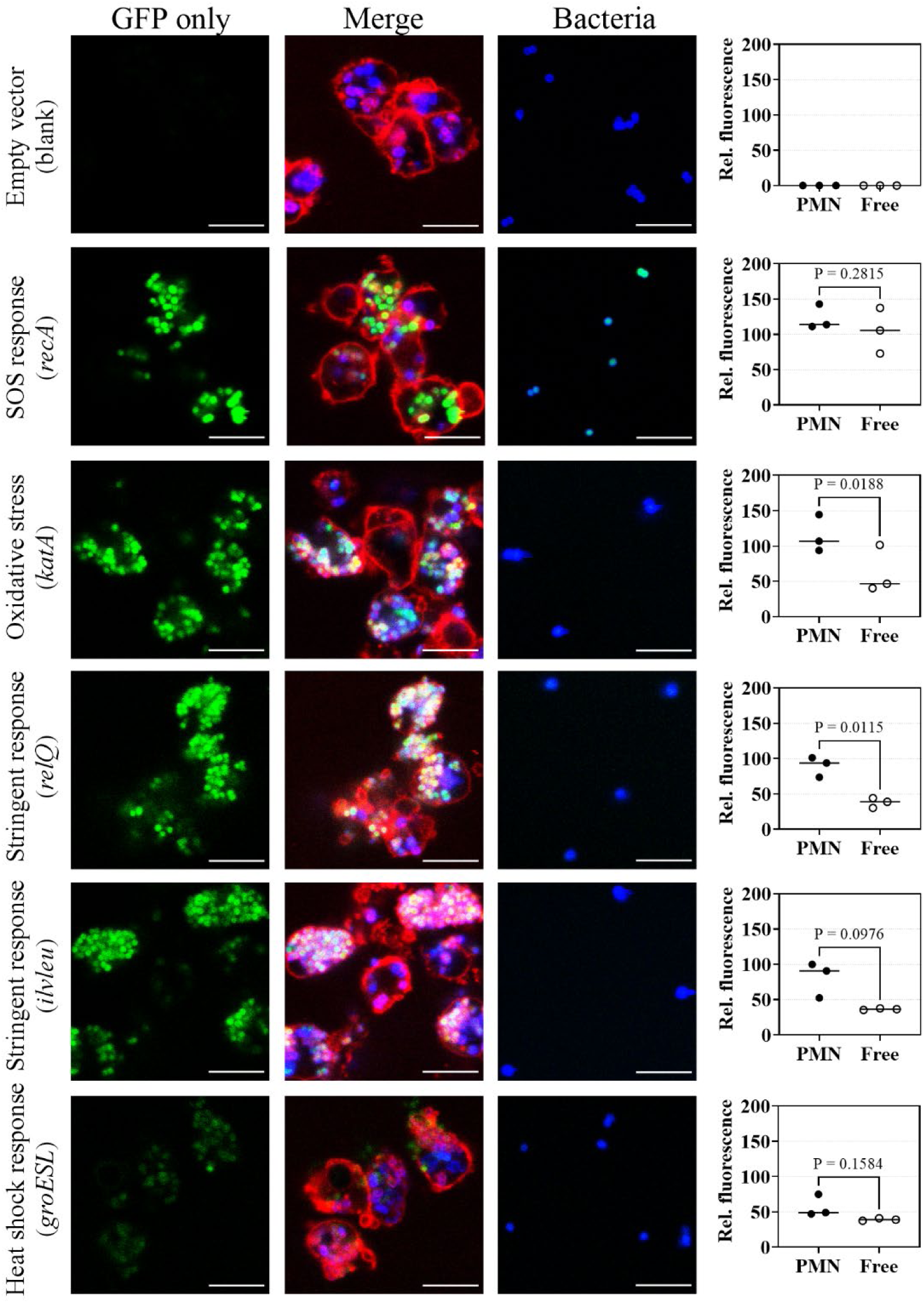
Activation of stress responses during phagocytosis in PMNs. Stress response reporter strains and the empty plasmid control strain were grown to mid-exponential phase in TSB supplemented with 0.1 mM HADA (counterstain). Subsequently, the reporter strains were mixed with PMNs in HBSS buffer or HBSS buffer alone (no PMN control) and human serum was added at a working concentration of 10% to initiate phagocytosis. PMN’s and bacteria were incubated for 2 h at 37 °C and 5% CO_2_ before washing resuspending in 2 mM EDTA. Samples were fixed in 4% PFA for 45 min before washing and resuspending in PBS + 3% BSA. Immediately before microscopy, neutrophils were stained with Alexa Fluor 594 conjugated anti-CD-16 antibodies at a concentration of 10 mg/L and samples were mounted on a microscope slide with antifade. Samples were imaged by CLSM using a 488 nm laser for GFP (green) excitation, a 405 nm laser for HADA (blue) excitation and a 555 nm laser to excite the anti-CD-16 antibodies (red). n = 3 biological replicates. Scale bar: 10 µm. The relative activation of stress responses were measured by GFP intensity of phagocytosed reporter strains (PMN) or reporter strains subjected to identical treatment, but without PMNs (free). Statistical testing was performed using a paired *t*-test.

Phagocytosis significantly activated the oxidative stress response and both branches of the stringent response, reported by *relQ* and *ilvleu* (Fig. 7). Activation of the heat shock reporter did not reach statistical significance. Among the phagocytosed bacteria, activation was again apparent across the population: cells within phagosomes were uniformly fluorescent rather than showing a mixture of bright and dark cells.

Activation of the SOS reporter also did not reach statistical significance, which was unexpected given the reactive oxygen species generated within the phagosome. The reason lies in the control rather than in the phagocytosed population. A small number of bacteria in the no-neutrophil control samples showed strong SOS reporter fluorescence, and this variability was sufficient to obscure any difference between the two groups, even though bacteria within phagosomes were consistently fluorescent. We therefore do not interpret this result as evidence that the SOS response is inactive during phagocytosis.

## Discussion

Using a set of fluorescent transcriptional reporters, we found that biofilms of *Staphylococcus aureus* grown in laboratory medium showed little or no activation of the SOS, oxidative stress, stringent or heat shock responses, whereas biofilms of the same strains grown in human serum activated all five reporters, in essentially all cells. Short exposure to serum activated both the cell wall stress-induced and the starvation-induced branches of the stringent response and abolished killing by ciprofloxacin. The comparison between biofilms grown in TSB and in serum holds the mode of growth, the substratum and the incubation period constant and varies only the chemical environment, and it indicates that the host environment rather than the biofilm growth mode is the operative variable.

### Biofilm phenotype does not induce stress in all laboratory biofilm models

The recalcitrance of biofilm infections to antibiotics is commonly explained by properties of the biofilm itself: mass transfer limitation generates gradients of oxygen and nutrients, and the resulting starvation and growth arrest render cells in the deeper layers refractory to killing. Nguyen et al. ^7^ established the general form of this argument, showing that active starvation responses rather than passive dormancy mediate tolerance in biofilms and in nutrient-limited bacteria. Based on this model, biofilm growth in a rich laboratory medium should be sufficient to activate the starvation-responsive arm of the stringent response. We did not observe this. Despite forming dense microcolonies over 24 h, biofilms grown in TSB showed only low-level reporter activity, close to that of the empty-plasmid control.

Two bodies of work bear on this. The first concerns how biofilms differ between laboratory and host-relevant conditions. Beenken et al. ^40^ characterised the global transcriptional profile of laboratory-grown *S. aureus* biofilms, and Cardile et al. ^41^ showed that human plasma enhances expression of adhesive matrix molecules, promotes biofilm formation and increases antimicrobial tolerance. Wieland et al. ^42^ reported divergence between *in vitro* and *in vivo* biofilm formation capacity in the same strains. Our observation extends this line by showing that the difference extends to bacterial stress physiology itself, not only to matrix composition and adhesion.

The second concerns whether laboratory biofilms are nutrient-restricted at all. Freiberg et al. ^43^ found arginine to be undetectable within *S. aureus* colony biofilms grown on both rich and chemically defined agar, and showed that restricting arginine induces tolerance to vancomycin, ceftaroline and delafloxacin through inhibition of protein synthesis, with the stringent response identified as the principal mechanism. Their biofilms were grown for 48 h at a solid–air interface, ours for 24 h submerged in excess broth, and the difference in outcome most plausibly reflects the degree of nutrient restriction each model imposes. The conclusion we draw is therefore not that laboratory biofilms are never stressed, but that whether they are depends strongly on the model, whereas the addition of serum produced strong activation in the very assay in which laboratory medium produced almost none. Biofilm growth mode is not sufficient in itself; the chemical environment determines the outcome.

The implication for experimental design is direct and is one we share with recent work from other groups. Ledger et al. ^44^ note that host-induced adaptations require exposure to host conditions and develop over time, and these features are consequently absent in *in vitro* biofilms grown in standard laboratory broth. Antimicrobial compounds screened against biofilms grown in laboratory media are screened against bacteria that may not be mounting the stress responses they mount during infection.

### Serum components activate stress

Heat inactivation of serum at 56 °C did not abolish reporter activation, indicating that complement proteins and other heat-labile serum enzymes are not required. This narrows the candidate space, and the leading candidates are all heat-stable.

The first is nutritional immunity. Free iron and zinc in serum are sequestered by transferrin and calprotectin ^45, 46^, and free amino acids are limited. This is consistent with activation of both stringent response branches, and with the arginine-dependent mechanism of Freiberg et al. ^43^, in which restriction of a single amino acid inhibits protein synthesis and triggers the stringent response. It may also explain why the oxidative stress reporter was activated in serum despite the absence of an exogenous oxidant. Fritsch et al. ^35^ showed that (p)ppGpp confers tolerance to oxidative stress by maintaining redox and iron homeostasis, which places metal limitation, the stringent response, and oxidative stress on a single axis rather than in separate compartments.

The second is host defence peptides. Ledger et al. ^22^ showed that LL-37 triggers the staphylococcal GraRS two-component system, leading to peptidoglycan accumulation and daptomycin tolerance, and Ledger and Edwards ^47^ subsequently showed that the resulting thickened cell wall conceals bound opsonins and impairs neutrophil killing. LL-37 is a small peptide and would survive our heat treatment.

The third, and most recently identified, is serum lipid. Ledger et al. ^44^ showed that the staphylococcal Geh lipase liberates glycerol from host triglycerides to support increased synthesis of D-alanylated wall teichoic acids, driving biofilm formation with accompanying vancomycin and daptomycin tolerance. Triglycerides are likewise heat-stable. Their design and ours are complementary rather than overlapping. They incubated bacteria in serum and then allowed biofilms to form in laboratory medium, whereas we grew biofilms in serum throughout. That two different arrangements of serum exposure and biofilm growth both produce tolerance suggests the effect is robust to the order in which they occur.

Our data do not identify which component activates the reporters, and several may act together. What the heat-inactivation result contributes is the exclusion of complement, otherwise among the most conspicuous candidates in serum.

### Serum-induced antibiotic tolerance is specific to ciprofloxacin

A 1 h exposure to serum abolished killing by ciprofloxacin but had little or no effect on killing by daptomycin, vancomycin, dicloxacillin or rifampicin. This appears at first to conflict with Ledger et al. ^22^, who found that serum triggered tolerance to daptomycin, vancomycin and several other agents. We suggest the difference is one of kinetics rather than of substance, and that two features of the literature account for it.

The first concerns ciprofloxacin. Fluoroquinolone killing depends on active DNA replication ^48^, and a substantial body of work implicates reactive oxygen species generated by active metabolism as a major contributor to quinolone-mediated death ^49–51^. Both requirements fail in a growth-arrested, metabolically quiescent cell. Tolerance to ciprofloxacin should therefore appear as soon as growth ceases, which is consistent with a 1 h serum exposure being sufficient.

The second concerns daptomycin. Ledger and Edwards ^52^ showed that daptomycin tolerance in growth-arrested *S. aureus* is not immediate: it develops over hours, requires glucose and active biosynthesis, and depends on peptidoglycan accumulation, failing to appear at 4 °C or without fermentable carbohydrate. They contrast this explicitly with tolerance to most other bactericidal antibiotics, which arises almost as soon as growth stops through the absence of replication and metabolic activity. Ledger et al. ^22^ used a 6 h serum exposure and Ledger et al. ^44^ a 16 h exposure, in both cases long enough for cell wall remodelling to occur, while our 1 h exposure is not.

Our 1 h result should therefore be read as capturing the earliest phase of host adaptation and as a lower bound on the effect of serum, rather than as a contradiction of the broader literature. Consistent with this, 1 h in serum activated only the two stringent response reporters, whereas 24 h of biofilm growth in serum activated all five. The emerging picture is of at least two temporally distinct routes to tolerance in the host: a rapid, largely passive one arising from growth arrest and loss of metabolic activity, protecting against agents that require active targets; and a slower, active one requiring biosynthesis and envelope remodelling, protecting against membrane- and cell wall-targeting agents. Our data address the first.

### Population-wide activation points to tolerance rather than persistence

Tolerance and persistence are distinguished by whether the surviving fraction comprises the whole population or a subpopulation ^2^, and persistence represents a dormant metabolic state that requires longer time for the cell to regain metabolic activity. Two features of our data indicate that what we observe is tolerance. Reporter activation in serum was apparent throughout the biofilms and in essentially all cells identified by the HADA counterstain, rather than in a scattered minority against an unlabelled background; and killing curves after serum exposure showed no killing at all, rather than the biphasic pattern characteristic of a susceptible majority with a surviving persister fraction.

This must be set against substantial evidence for heterogeneity in this organism. Korshoj and Kielian ^53^, using single-cell RNA sequencing, resolved seven transcriptional subpopulations within *S. aureus* biofilms, including a transcriptionally dormant cluster, and found responses tailored to the immune cell type encountered. Hira et al. ^54^ identified dormant and bimodally growing subpopulations within isogenic communities and showed these are not explained by position within a colony. Huemer et al. ^9^ recovered growth-arrested, antibiotic-tolerant bacteria from patient material, where the tolerant phenotype was confined to a fraction of the population.

We do not regard these findings as incompatible with ours. The first two were obtained from biofilms and colonies grown in laboratory media, which our data indicate is a physiologically distinct state from growth in serum. More fundamentally, heterogeneity across an entire transcriptome, in carbon metabolism and virulence regulons, does not entail heterogeneity at a small number of stress response promoters: a population may be transcriptionally diverse in general while responding uniformly to a strong and uniformly distributed environmental signal. It is also relevant that Korshoj and Kielian ^53^ identified the construction of fluorescent reporters for genes enriched in distinct transcriptional clusters as a needed next step, which is the kind of application the strains described here are intended to support.

### Linking stress responses and growth arrest to antibiotic tolerance

Activation of either branch of the stringent response, or of the heat shock response, reduced the rate of ciprofloxacin killing. Each of these pre-treatments also arrested growth for the duration of the antibiotic challenge, so the two variables cannot be separated in our experiments, and we describe the association as one between growth-arresting stress conditions and reduced killing.

There is good reason to think this reflects the underlying biology rather than only a limitation of design. Conlon et al. ^5^ found that a mutant unable to synthesise (p)ppGpp remained as susceptible to ciprofloxacin as the wild type, and proposed ATP depletion rather than alarmone signalling as the proximate cause. Kwan et al. ^8^ showed that arrested protein synthesis alone increases the formation of persister-like cells. Ledger and Edwards ^52^ found that growth arrest imposed with any of several unrelated antibiotics was sufficient to induce tolerance. Freiberg et al. ^43^ found that restricting a single amino acid, or inhibiting protein synthesis by any of three pharmacological routes, produced tolerance in biofilms. Huemer et al. ^55^ identified serine-threonine phosphoregulation by PknB and Stp as a further route into a quiescent, tolerant state. In each case a different upstream signal converges on the same downstream physiology.

The consistent reading is that the arrest of growth and biosynthesis is the proximate determinant of tolerance to antibiotics whose activity depends on active targets, and that the stringent response is one of several routes into that state rather than the mediator of tolerance as such. This does not diminish the role of the stringent response in survival. Geiger et al. ^18^ showed that RSH-dependent (p)ppGpp synthesis is required for full virulence, and Geiger et al. ^56^ showed that the small synthetases RelP and RelQ mediate tolerance to cell envelope stress, which is consistent with our observation that vancomycin pre-treatment activated the *relQ* reporter and slowed ciprofloxacin killing. It does, however, mean that our data cannot attribute tolerance to stringent response signalling in isolation. The point bears on the serum result as well, since *S. aureus* does not replicate in human serum ^44^. The growth arrest imposed by the host environment may be sufficient to account for the loss of ciprofloxacin killing, with stress response activation as a correlate of that state.

### Stress responses during phagocytosis

Inside human neutrophils, the reporters for the oxidative stress response and both the cell wall stress-induced and the starvation-induced branches of the stringent response were significantly activated. This corroborates at single-cell resolution what previous work inferred from bulk measurements. Voyich et al. ^57^ documented broad transcriptional reprogramming of *S. aureus* after phagocytosis, including SOS, stringent, cell wall stress and heat shock genes, and Peyrusson et al. ^12^ found a comparable signature in intracellular bacteria surviving antibiotic exposure. Peyrusson et al. ^15^ and Rowe et al. ^16^ have both linked host-derived oxidative stress to the tolerant intracellular phenotype, and Geiger et al. ^56^ showed that loss of RSH-dependent (p)ppGpp synthesis reduces survival after phagocytosis, connecting the responses we observe to a measurable survival benefit. Activation of the heat shock reporter did not reach statistical significance, and the SOS comparison was confounded by variability among bacteria in the neutrophil-free control.

The clinical significance of this compartment is well established. Most antibiotics penetrate neutrophils poorly or lose activity in the phagosomal environment ^58^, so intracellular bacteria are shielded both by their location and, our data suggest, by their physiological state. One design consideration applies. Ledger and Edwards ^47^ showed that prolonged serum incubation thickens the cell wall and conceals bound opsonins, reducing phagocytosis and killing by neutrophils. Our opsonisation used 10% serum briefly, a condition that did not activate the reporters in the biofilm experiments, so extensive remodelling is unlikely; nonetheless, any phagocytosis assay includes a serum exposure component, and the two host signals cannot be entirely separated.

### Limitations

Several constraints bound this work. Each stress response is represented by a single promoter. For example, *katA* reports only one arm of a regulon that also encompasses superoxide dismutase and the PerR-controlled iron storage proteins ^14, 25^, and *groESL* reports the accumulation of misfolded protein rather than temperature as such, with the Clp chaperones and proteases constituting a further arm of that response ^38^. The relationship between stress response activation and tolerance is correlative and only tested in a single strain. We did not test mutants defective in individual pathways.

### Implications

Our study has two major implications. Firstly, our results show that omitting host components from in vitro biofilm models may substantially misrepresent the physiological state of bacteria during infection. Including specific host components could make such models more predictive of therapeutic outcome. Secondly, our results suggest that arrest of growth and biosynthesis is the proximate cause of tolerance to replication-dependent antibiotics. Novel interventions could be directed at preventing bacteria from entering or maintaining that state. Recent work identifying the Geh lipase and wall teichoic acid synthesis as tractable targets in serum-adapted bacteria ^44^, and fosfomycin as an inhibitor of host-induced cell wall thickening that restores opsonophagocytic killing ^47^, illustrates the value of targeting the host-adaptation process itself. Screening approaches directed at non-growing populations, such as that described by Petersen et al. ^6^, are complementary.

The reporter strains described here are deposited at Addgene and can be transferred to other genetic backgrounds, including methicillin-resistant clinical isolates. They provide a means of asking, in situ and cell by cell, which stress responses bacteria mount in environments that cannot be sampled by bulk methods.

## Acknowledgements

This work was funded by the Novo Nordisk Foundation (grant no. NNF19OC0058357)

## References

(1) Brauner, A.; Fridman, O.; Gefen, O.; Balaban, N. Q. Distinguishing between resistance, tolerance and persistence to antibiotic treatment. Nature Publishing Group: 2016; Vol. 14, pp 320–330.

(2) Balaban, N. Q.; Helaine, S.; Lewis, K.; Ackermann, M.; Aldridge, B.; Andersson, D. I.; Brynildsen, M. P.; Bumann, D.; Camilli, A.; Collins, J. J.;, et al. Definitions and guidelines for research on antibiotic persistence. Nature Reviews Microbiology 2019, 17 (7), 441–448. DOI: 10.1038/s41579-019-0196-3.

(3) Young, M. H.; Upchurch, G. R.; Malani, P. N. Vascular Graft Infections. Elsevier: 2012; Vol. 26, pp 41–56.

(4) Tong, S. Y. C.; Davis, J. S.; Eichenberger, E.; Holland, T. L.; Fowler, V. G. Staphylococcus aureus infections: Epidemiology, pathophysiology, clinical manifestations, and management. Clinical Microbiology Reviews 2015, 28 (3), 603–661. DOI: 10.1128/CMR.00134-14.

(5) Conlon, B. P.; Rowe, S. E.; Gandt, A. B.; Nuxoll, A. S.; Donegan, N. P.; Zalis, E. A.; Clair, G.; Adkins, J. N.; Cheung, A. L.; Lewis, K. Persister formation in Staphylococcus aureus is associated with ATP depletion. Nature Microbiology 2016, 1 (5). DOI: 10.1038/nmicrobiol.2016.51.

(6) Petersen, M. E.; Hansen, L. K.; Mitkin, A. A.; Kelly, N. M.; Wood, T. K.; Jørgensen, N. P.; Østergaard, L. J.; Meyer, R. L. A high-throughput assay identifies molecules with antimicrobial activity against persister cells. Journal of Medical Microbiology 2024, 73 (7), 1–12. DOI: 10.1099/jmm.0.001856.

(7) Nguyen, D.; Joshi-Datar, A.; Lepine, F.; Bauerle, E.; Olakanmi, O.; Beer, K.; McKay, G.; Siehnel, R.; Schafhauser, J.; Wang, Y.;, et al. Active starvation responses mediate antibiotic tolerance in biofilms and nutrient-limited bacteria. Science 2011, 334 (6058), 982–986. DOI: 10.1126/science.1211037.

(8) Kwan, B. W.; Valenta, J. A.; Benedik, M. J.; Wood, T. K. Arrested protein synthesis increases persister-like cell formation. Antimicrobial Agents and Chemotherapy 2013, 57 (3), 1468–1473. DOI: 10.1128/AAC.02135-12.

(9) Huemer, M.; Shambat, S. M.; Bergada-Pijuan, J.; Söderholm, S.; Boumasmoud, M.; Vulin, C.; Gómez-Mejia, A.; Varela, M. A.; Tripathi, V.; Götschi, S.;, et al. Molecular reprogramming and phenotype switching in Staphylococcus aureus lead to high antibiotic persistence and affect therapy success. Proceedings of the National Academy of Sciences of the United States of America 2021, 118 (7), 1–12. DOI: 10.1073/pnas.2014920118.

(10) Lacoma, A.; Edwards, A. M.; Young, B. C.; Domínguez, J.; Prat, C.; Laabei, M. Cigarette smoke exposure redirects Staphylococcus aureus to a virulence profile associated with persistent infection. Scientific Reports 2019, 9 (1), 1–15. DOI: 10.1038/s41598-019-47258-6.

(11) Painter, K. L.; Strange, E.; Parkhill, J.; Bamford, K. B.; Armstrong-James, D.; Edwards, A. M. Staphylococcus aureus adapts to oxidative stress by producing H2O2-resistant small-colony variants via the SOS response. Infection and Immunity 2015, 83 (5), 1830–1844. DOI: 10.1128/IAI.03016-14.

(12) Peyrusson, F.; Varet, H.; Nguyen, T. K.; Legendre, R.; Sismeiro, O.; Coppée, J. Y.; Wolz, C.; Tenson, T.; Van Bambeke, F. Intracellular Staphylococcus aureus persisters upon antibiotic exposure. Nature Communications 2020, 11 (1). DOI: 10.1038/s41467-020-15966-7.

(13) Vestergaard, M.; Paulander, W.; Ingmer, H. Activation of the SOS response increases the frequency of small colony variants. BMC Research Notes 2015, 8 (1), 1–5. DOI: 10.1186/s13104-015-1735-2.

(14) Horsburgh, M. J.; Clements, M. O.; Crossley, H.; Ingham, E.; Foster, S. J. PerR controls oxidative stress resistance and iron storage proteins and is required for virulence in Staphylococcus aureus. Infection and Immunity 2001, 69 (6), 3744–3754. DOI: 10.1128/IAI.69.6.3744-3754.2001.

(15) Peyrusson, F.; Nguyen, T. K.; Najdovski, T.; Van Bambeke, F. Host Cell Oxidative Stress Induces Dormant Staphylococcus aureus Persisters. Microbiology Spectrum 2022, 10 (1). DOI: 10.1128/spectrum.02313-21.

(16) Rowe, S. E.; Wagner, N. J.; Li, L.; Beam, J. E.; Wilkinson, A. D.; Radlinski, L. C.; Zhang, Q.; Miao, E. A.; Conlon, B. P. Reactive oxygen species induce antibiotic tolerance during systemic Staphylococcus aureus infection. Nature Microbiology 2020, 5 (2), 282–290. DOI: 10.1038/s41564-019-0627-y.

(17) Berti, A. D.; Shukla, N.; Rottier, A. D.; McCrone, J. S.; Turner, H. M.; Monk, I. R.; Baines, S. L.; Howden, B. P.; Proctor, R. A.; Rose, W. E. Daptomycin selects for genetic and phenotypic adaptations leading to antibiotic tolerance in MRSA. Journal of Antimicrobial Chemotherapy 2018, 73 (8), 2030–2033. DOI: 10.1093/jac/dky148.

(18) Geiger, T.; Goerke, C.; Fritz, M.; Schäfer, T.; Ohlsen, K.; Liebeke, M.; Lalk, M.; Wolz, C. Role of the (p)ppGpp synthase RSH, a RelA/SpoT homolog, in stringent response and virulence of Staphylococcus aureus. Infection and Immunity 2010, 78 (5), 1873–1883. DOI: 10.1128/IAI.01439-09.

(19) Geiger, T.; Kästle, B.; Gratani, F. L.; Goerke, C.; Wolz, C. Two small (p)ppGpp synthases in staphylococcus aureus mediate tolerance against cell envelope stress conditions. Journal of Bacteriology 2014, 196 (4), 894–902. DOI: 10.1128/JB.01201-13.

(20) Matsuo, M.; Hiramatsu, M.; Singh, M.; Sasaki, T.; Hishinuma, T.; Yamamoto, N.; Morimoto, Y.; Kirikae, T.; Hiramatsu, K. Genetic and transcriptomic analyses of ciprofloxacin-tolerant staphylococcus aureus isolated by the replica plating tolerance isolation system (REPTIS). Antimicrobial Agents and Chemotherapy 2019, 63 (2). DOI: 10.1128/AAC.02019-18.

(21) Singh, M.; Matsuo, M.; Sasaki, T.; Morimoto, Y.; Hishinuma, T.; Hiramatsu, K. In Vitro tolerance of drug-naive Staphylococcus aureus strain FDA209P to vancomycin. Antimicrobial Agents and Chemotherapy 2017, 61 (2). DOI: 10.1128/AAC.01154-16.

(22) Ledger, E. V. K.; Mesnage, S.; Edwards, A. M. Human serum triggers antibiotic tolerance in Staphylococcus aureus. Nature Communications 2022, 13 (1), 1–19. DOI: 10.1038/s41467-022-29717-3.

(23) Crameri, A.; Whitehorn, E. A.; Tate, E.; Stemmer, W. P. Improved green fluorescent protein by molecular evolution using DNA shuffling. Nat Biotechnol 1996, 14 (3), 315–319. DOI: 10.1038/nbt0396-315 From NLM.

(24) Singh, V. K.; Syring, M.; Singh, A.; Singhal, K.; Dalecki, A.; Johansson, T. An insight into the significance of the DnaK heat shock system in Staphylococcus aureus. International Journal of Medical Microbiology 2012, 302 (6), 242–252. DOI: 10.1016/j.ijmm.2012.05.001.

(25) Cheng, K.; Sun, Y.; Yu, H.; Hu, Y.; He, Y.; Shen, Y. Staphylococcus aureus SOS response: Activation, impact, and drug targets. mLife 2024, 3 (3), 343–366. DOI: 10.1002/mlf2.12137.

(26) Úbeda, C.; Maiques, E.; Knecht, E.; Lasa, Í.; Novick, R. P.; Penadés, J. R. Antibiotic-induced SOS response promotes horizontal dissemination of pathogenicity island-encoded virulence factors in staphylococci. Molecular Microbiology 2005, 56 (3), 836–844. DOI: 10.1111/j.1365-2958.2005.04584.x.

(27) Maiques, E.; Úbeda, C.; Campoy, S.; Salvador, N.; Lasa, Í.; Novick, R. P.; Barbé, J.; Penadés, J. R. β-lactam antibiotics induce the SOS response and horizontal transfer of virulence factors in Staphylococcus aureus. Journal of Bacteriology 2006, 188 (7), 2726–2729. DOI: 10.1128/JB.188.7.2726-2729.2006.

(28) Chang, W.; Small, D. A.; Toghrol, F.; Bentley, W. E. Global transcriptome analysis of Staphylococcus aureus response to hydrogen peroxide. Journal of Bacteriology 2006, 188 (4), 1648–1659. DOI: 10.1128/jb.188.4.1648-1659.2006.

(29) Anderson, K. L.; Roberts, C.; Disz, T.; Vonstein, V.; Hwang, K.; Overbeek, R.; Olson, P. D.; Projan, S. J.; Dunman, P. M. Characterization of the Staphylococcus aureus heat shock, cold shock, stringent, and SOS responses and their effects on log-phase mRNA turnover. Journal of Bacteriology 2006, 188 (19), 6739– 6756. DOI: 10.1128/JB.00609-06.

(30) Ha, K. P.; Edwards, A. M. DNA Repair in Staphylococcus aureus. Microbiology and Molecular Biology Reviews 2021, 85 (4). DOI: 10.1128/mmbr.00091-21.

(31) Cirz, R. T.; Jones, M. B.; Gingles, N. A.; Minogue, T. D.; Jarrahi, B.; Peterson, S. N.; Romesberg, F. E. Complete and SOS-mediated response of Staphylococcus aureus to the antibiotic ciprofloxacin. Journal of Bacteriology 2007, 189 (2), 531–539. DOI: 10.1128/JB.01464-06.

(32) Clements, M. O.; Watson, S. P.; Foster, S. J. Characterization of the major superoxide dismutase of Staphylococcus aureus and its role in starvation survival, stress resistance, and pathogenicity. Journal of Bacteriology 1999, 181 (13), 3898–3903. DOI: 10.1128/jb.181.13.3898-3903.1999.

(33) Bore, E.; Langsrud, S.; Langsrud, Ø.; Rode, T. M.; Holck, A. Acid-shock responses in Staphylococcus aureus investigated by global gene expression analysis. Microbiology 2007, 153 (7), 2289–2303. DOI: 10.1099/mic.0.2007/005942-0.

(34) Irving, S. E.; Choudhury, N. R.; Corrigan, R. M. The stringent response and physiological roles of (pp)pGpp in bacteria. 2021; Vol. 19, pp 256–271.

(35) Fritsch, V. N.; Loi, V. V.; Busche, T.; Tung, Q. N.; Lill, R.; Horvatek, P.; Wolz, C.; Kalinowski, J.; Antelmann, H. The alarmone (p)ppGpp confers tolerance to oxidative stress during the stationary phase by maintenance of redox and iron homeostasis in Staphylococcus aureus. Free Radical Biology and Medicine 2020, 161, 351–364. DOI: 10.1016/j.freeradbiomed.2020.10.322.

(36) Laport, M. S.; De Castro, A. C. D.; Villardo, A.; Lemos, J. A. C.; Bastos, M. D. C. F.; Giambiagi-deMarval, M. Expression of the major heat shock proteins dnaK and groEL in Streptococcus pyogenes: A comparison to Enterococcus faecalis and Staphylococcus aureus. Current Microbiology 2001, 42 (4), 264–268. DOI: 10.1007/s002840110215.

(37) Lund, P. A. Multiple chaperonins in bacteria - Why so many?: Review article. FEMS Microbiology Reviews 2009, 33 (4), 785–800. DOI: 10.1111/j.1574-6976.2009.00178.x.

(38) Frees, D.; Gerth, U.; Ingmer, H. Clp chaperones and proteases are central in stress survival, virulence and antibiotic resistance of Staphylococcus aureus. International Journal of Medical Microbiology 2014, 304 (2), 142–149. DOI: 10.1016/j.ijmm.2013.11.009.

(39) Monk, I. R.; Shah, I. M.; Xu, M.; Tan, M. W.; Foster, T. J. Transforming the untransformable: Application of direct transformation to manipulate genetically Staphylococcus aureus and Staphylococcus epidermidis. mBio 2012, 3 (2). DOI: 10.1128/mBio.00277-11.

(40) Beenken, K. E.; Dunman, P. M.; McAleese, F.; Macapagal, D.; Murphy, E.; Projan, S. J.; Blevins, J. S.; Smeltzer, M. S. Global gene expression in Staphylococcus aureus biofilms. J Bacteriol 2004, 186 (14), 4665– 4684. DOI: 10.1128/jb.186.14.4665-4684.2004 From NLM.

(41) Cardile, A. P.; Sanchez, C. J., Jr.; Samberg, M. E.; Romano, D. R.; Hardy, S. K.; Wenke, J. C.; Murray, C. K.; Akers, K. S. Human plasma enhances the expression of Staphylococcal microbial surface components recognizing adhesive matrix molecules promoting biofilm formation and increases antimicrobial tolerance In Vitro. BMC Res Notes 2014, 7, 457. DOI: 10.1186/1756-0500-7-457 From NLM.

(42) Wieland, B.; Gunaratnam, G.; Pätzold, L.; Wadood, N. A.; Schmartz, G. P.; Kundu, S.; Kirilov, N. K.; Krüger, I.; Elhawy, M. I.; Rehner, J.;, et al. Assessment of the biofilm formation capacities of Staphylococcus aureus strains Newman and Newman D2C in vitro and in vivo. Scientific Reports 2025, 15 (1), 16132. DOI: 10.1038/s41598-025-00521-5.

(43) Freiberg, J. A.; Reyes Ruiz, V. M.; Gimza, B. D.; Murdoch, C. C.; Green, E. R.; Curry, J. M.; Cassat, J. E.; Skaar, E. P. Restriction of arginine induces antibiotic tolerance in Staphylococcus aureus. Nature Communications 2024, 15 (1), 6734. DOI: 10.1038/s41467-024-51144-9.

(44) Ledger, E. V. K.; Horgan, N. E.; Lynch, D.; Bateman, L. M.; Massey, R. C. Human serum triglycerides promote Staphylococcus aureus biofilm formation and antibiotic tolerance. npj Biofilms and Microbiomes 2026. DOI: 10.1038/s41522-026-01064-x.

(45) Hood, M. I.; Skaar, E. P. Nutritional immunity: transition metals at the pathogen-host interface. Nat Rev Microbiol 2012, 10 (8), 525–537. DOI: 10.1038/nrmicro2836 From NLM.

(46) Zygiel, E. M.; Nolan, E. M. Transition Metal Sequestration by the Host-Defense Protein Calprotectin. Annu Rev Biochem 2018, 87, 621–643. DOI: 10.1146/annurev-biochem-062917-012312 From NLM.

(47) Ledger, E. V. K.; Edwards, A. M. Host-induced cell wall remodeling impairs opsonophagocytosis of Staphylococcus aureus by neutrophils. mBio 2024, 15 (8), e0164324. DOI: 10.1128/mbio.01643-24 From NLM.

(48) Gradelski, E.; Kolek, B.; Bonner, D.; Fung-Tomc, J. Bactericidal mechanism of gatifloxacin compared with other quinolones. Journal of Antimicrobial Chemotherapy 2002, 49 (1), 185–188. DOI: 10.1093/jac/49.1.185.

(49) Hong, Y.; Li, Q.; Gao, Q.; Xie, J.; Huang, H.; Drlica, K.; Zhao, X. Reactive oxygen species play a dominant role in all pathways of rapid quinolone-mediated killing. Journal of Antimicrobial Chemotherapy 2020, 75 (3), 576–585. DOI: 10.1093/jac/dkz485.

(50) Kottur, J.; Nair, D. T. Reactive oxygen species play an important role in the bactericidal activity of quinolone antibiotics. Angewandte Chemie - International Edition 2016, 55 (7), 2397–2400. DOI: 10.1002/anie.201509340.

(51) Wang, X.; Zhao, X.; Malik, M.; Drlica, K. Contribution of reactive oxygen species to pathways of quinolone-mediated bacterial cell death. Journal of Antimicrobial Chemotherapy 2010, 65 (3), 520–524. DOI: 10.1093/jac/dkp486.

(52) Ledger, E. V. K.; Edwards, A. M. Growth Arrest of Staphylococcus aureus Induces Daptomycin Tolerance via Cell Wall Remodelling. mBio 2023, 14 (1), e0355822. DOI: 10.1128/mbio.03558-22 From NLM.

(53) Korshoj, L. E.; Kielian, T. Bacterial single-cell RNA sequencing captures biofilm transcriptional heterogeneity and differential responses to immune pressure. Nature Communications 2024, 15 (1), 10184. DOI: 10.1038/s41467-024-54581-8.

(54) Hira, J.; Singh, B.; Halder, T.; Mahmutovic, A.; Ajayi, C.; Sekh, A. A.; Hegstad, K.; Johannessen, M.; Lentz, C. S. Single-cell phenotypic profiling and backtracing exposes and predicts clinically relevant subpopulations in isogenic Staphylococcus aureus communities. Communications Biology 2024, 7 (1), 1228. DOI: 10.1038/s42003-024-06894-z.

(55) Huemer, M.; Mairpady Shambat, S.; Hertegonne, S.; Bergada-Pijuan, J.; Chang, C. C.; Pereira, S.; Gómez-Mejia, A.; Van Gestel, L.; Bär, J.; Vulin, C.;, et al. Serine-threonine phosphoregulation by PknB and Stp contributes to quiescence and antibiotic tolerance in Staphylococcus aureus. Science signaling 2023, 16 (766), eabj8194–eabj8194. DOI: 10.1126/scisignal.abj8194.

(56) Geiger, T.; Francois, P.; Liebeke, M.; Fraunholz, M.; Goerke, C.; Krismer, B.; Schrenzel, J.; Lalk, M.; Wolz, C. The Stringent Response of Staphylococcus aureus and Its Impact on Survival after Phagocytosis through the Induction of Intracellular PSMs Expression. PLoS Pathogens 2012, 8 (11), e1003016–e1003016. DOI: 10.1371/journal.ppat.1003016.

(57) Voyich, J. M.; Braughton, K. R.; Sturdevant, D. E.; Whitney, A. R.; Porcella, S. F.; Long, R. D.; Dorward, D. W.; Gardner, D. J.; Kreiswirth, B. N.; Musser, J. M.;, et al. Insights into Mechanisms Used by Staphylococcus aureus to Avoid Destruction by Human Neutrophils. The Journal of Immunology 2005, 175 (2005), 3907– 3919.

(58) Bongers, S.; Hellebrekers, P.; Leenen, L. P. H.; Koenderman, L.; Hietbrink, F. Intracellular penetration and effects of antibiotics on staphylococcus aureus inside human neutrophils: A comprehensive review. Antibiotics 2019, 8 (2), 1–22. DOI: 10.3390/antibiotics8020054.

