## Supplementary materials for "The host environment rather than biofilm growth activates stress responses and conveys ciprofloxacin tolerance in *Staphylococcus aureus*"

Table S1: Promotor sequences

Color coding:

Linker/spacer; Restriction site ; Assumed promotor region (394-500 bp upstream) ; Predicted promoter sequence ; Assumed Shine Dalgarno sequence ; Start codon ; $gfp_{uvr}$ gene (optimized to *S. aureus*) ; Stop codon ; Linker/spacer

| Gene fusion | Sequence |
| --- | --- |
| $recA-gfp_{uvr}$ | GAGCTCCGATGTTCTAAAGGGTATGATTAATCACAATGAAAACTTTGTTGATATTAATAAACCTATTGA<br>GCAGCAATTAAAAGATGCAGTGCAATTTGTTAATAAATTGTTTAATGTGTCATCAGCAATTATTCTATT<br>AGAGTATGATGGTGTAGTCCATATAGGCTATGATAATAACTTTGAATTTAAAACTGAGCAATTTAAAAT<br>GTCTAAATCTAGAAATTTATTAAAGAACAGAAGTCAAAATTATGTGCTCATAAGATTATTAAATTGGCT<br>TAGAACAACAAATTAATTGTATTATCGATAAAAATATAAGCACGTTTGTTCGTTTTTGCGTTTTGATTT<br>CAAGATTTTATACGAACAAATATTCGCAAAACACTTGTATTTTATTTTGAATCCTTGTATAGTATTGAT<br>AAGATAATTAAAAGATAGCAATTTCAATT**AGGAGG**TCTCGCTatgtcaaaggtgaagaattatttaca<br>ggtgtagtaccaattttagtagaattagatggtgatgtaaatggtcataaattttcagtatcaggtgaa<br>ggtgaaggtgatgcaacatatggtaaattaacattaaaatttatttgtacaacaggtaaattaccagta<br>ccatggccaacattagtaacaacatttacatatggtgtacaatgtttttcaagatatccagatcatatg<br>aaaagacatgatttttttaaatcagcaatgccagaaggttatgtacaagaaagaacaatttcatttaaa<br>gatgatggtaattataaaacaagagcagaagtaaaatttgaaggtgatacattagtaaatagaattgaa<br>ttaaaaggtattgattttaaagaagatggtaatattttaggtcataaattagaatataattataattca<br>cataatgtatatattacagcagataaacaaaaaaatggtattaaagcaaattttaaaattagacataat<br>attgaagatggttcagtacaattagcagatcattatcaacaaaatacaccaattggtgatggtccagta<br>ttattaccagataatcattatttatcaacacaatcagcattatcaaaagatccaaatgaaaaaagagat<br>catatggtattattagaatttgtaacagcagcaggtattacacatggtatggatgaattatataaataa<br>TAATTGAGCTCAACCT |
| $katA-gfp_{uvr}$ | GAGCTCTAACAAGATAAGCGAGTATAGCGCCTCCAGGACCAGCTTGAGAAATGATATTACCAGTAGCTA<br>CAAATAGACCAGTCCCAATTGCACCACCTATAGCAATCATGGAAATGTGTCTTGAGTTAAGACTACGGT<br>TCATTTTATTATCTTCCATATTTAGTCTCCCATCTATTTAAATATACCCATTATTGTAAGCTTTTTAAG<br>TGTACTATTCAATAACTATTTAGTACTGTAAAGCGAAAAAATTAAAATTTTCTGATTTTTTAATCATCT<br>TGAGCATGTTTAATTGTAATTTTGATGGGGTTAAATTATAATATGTATTAAATTATAATTATTATAAAT<br>TGTGGAGGGATGACTATGTCACAACAAGACAAAAAGTTAACTGGTGTTTTTGGGCATCCAGTATCAGAC<br>CGAGAAAATAGTATGACAGCAGGGCCT**AGGGGA**CCTCTTTTAatgtcaaaggtgaagaattatttaca<br>ggtgtagtaccaattttagtagaattagatggtgatgtaaatggtcataaattttcagtatcaggtgaa<br>ggtgaaggtgatgcaacatatggtaaattaacattaaaatttatttgtacaacaggtaaattaccagta<br>ccatggccaacattagtaacaacatttacatatggtgtacaatgtttttcaagatatccagatcatatg<br>aaaagacatgatttttttaaatcagcaatgccagaaggttatgtacaagaaagaacaatttcatttaaa<br>gatgatggtaattataaaacaagagcagaagtaaaatttgaaggtgatacattagtaaatagaattgaa<br>ttaaaaggtattgattttaaagaagatggtaatattttaggtcataaattagaatataattataattca<br>cataatgtatatattacagcagataaacaaaaaaatggtattaaagcaaattttaaaattagacataat<br>attgaagatggttcagtacaattagcagatcattatcaacaaaatacaccaattggtgatggtccagta<br>ttattaccagataatcattatttatcaacacaatcagcattatcaaaagatccaaatgaaaaaagagat<br>catatggtattattagaatttgtaacagcagcaggtattacacatggtatggatgaattatataaataa<br>TAATTGAGCTCAACCT |
| $relQ-gfp_{uvr}$ | GAGCTCGAACTTCGTACAATTCCATACAGCAACAATGGAAAAAACAGAGGTGGATAGTGAATATGCAAC<br>AACTGATAATGAAGAACATATTCAAGAACAATCAAATCAAGATAAATCGTACTTTGATAGCGAATCAAT<br>TGGCAACCAAAGTGGTGGCGTTCGACAAAGAGAACCGCTTATGGATGGTCCACATCCAAAAGTTGTGGA<br>ATCATTTAAAGACGTATAACATCTCTATCAATTAAGCACTTGTACCTATATAACGGTAACAGGTGTTTT<br>TTTGCGTTAAATCATTTTAAAATCACTTCCTGTAATTGAAGGTTTTGATACATAATTCGAATTGTTTGA<br>GTTTCTTTGTGCTGTTTCAGTATCAGCCCAAAGTTCGATAGTGTATTAGTATTATCTTAATAAAATGTT<br>AGGTACAATAAAGATGATTATATAT**CGGAGG**TTAGTATAAAAatgtcaaaggtgaagaattatttaca<br>ggtgtagtaccaattttagtagaattagatggtgatgtaaatggtcataaattttcagtatcaggtgaa<br>ggtgaaggtgatgcaacatatggtaaattaacattaaaatttatttgtacaacaggtaaattaccagta |

| | |
|---|---|
| | ccatggccaacattagtaacaacatttacatatggtgtacaatgtttttcaagatatccagatcatatg<br>aaaagacatgatttttttaaatcagcaatgccagaaggttatgtacaagaaagaacaatttcatttaaa<br>gatgatggtaattataaaacaagagcagaagtaaaatttgaaggtgatacattagtaaatagaattgaa<br>ttaaaaggtattgattttaaagaagatggtaatattttaggtcataaattagaatataattataattca<br>cataatgtatatattacagcagataaacaaaaaaatggtattaaagcaaattttaaaattagacataat<br>attgaagatggttcagtacaattagcagatcattatcaacaaaatacaccaattggtgatggtccagta<br>ttattaccagataatcattatttatcaacacaatcagcattatcaaaagatccaaatgaaaaaagagat<br>catatggtattattagaatttgtaacagcagcaggtattacacatggtatggatgaattatataaataa<br>TAATTGAGCTCAACCT |
| $groESL-gfp_{uvr}$ | GAGCTCGTGCTAAACTTTAGGTTTTTTAAGGAGGAACAATCATGCTAAAACCAATTGGAAATCGTGTGA<br>TTATTGAGAAAAAAGAACAAGAACAAACAACTAAAAGTGGTATTGTTTTAACTGATAGTGCTAAAGAAA<br>AATCAAACGAAGGCGTTATCGTTGCAGTAGGAACTGGACGTCTATTAAATGATGGTACAAGAGTGACTC<br>CTGAAGTGAAAGAAGGGGACCGTGTCGTGTTCCAACAATATGCTGGTACAGAAGTTAAACGAGATAATG<br>AAACATATCTAGTATTAAATGAAGAAGATATTTTAGCGGTAATTGAATAATATAAAATTAAATTCATAG<br>ATAAATTGTAAAGAACGAAAATGAAATATGACTAAACAAAT**GGAGG**TTTATCATTTatgtcaaaaggtg<br>aagaattatttacaggtgtagtaccaattttagtagaattagatggtgatgtaaatggtcataaatttt<br>cagtatcaggtgaaggtgaaggtgatgcaacatatggtaaattaacattaaaatttatttgtacaacag<br>gtaaattaccagtaccatggccaacattagtaacaacatttacatatggtgtacaatgtttttcaagat<br>atccagatcatatgaaaagacatgatttttttaaatcagcaatgccagaaggttatgtacaagaaagaa<br>caatttcatttaaagatgatggtaattataaaacaagagcagaagtaaaatttgaaggtgatacattag<br>taaatagaattgaattaaaaggtattgattttaaagaagatggtaatattttaggtcataaattagaat<br>ataattataattcacataatgtatatattacagcagataaacaaaaaaatggtattaaagcaaatttta<br>aaattagacataatattgaagatggttcagtacaattagcagatcattatcaacaaaatacaccaattg<br>gtgatggtccagtattattaccagataatcattatttatcaacacaatcagcattatcaaaagatccaa<br>atgaaaaaagagatcatatggtattattagaatttgtaacagcagcaggtattacacatggtatggatg<br>aattatataaataaTAATTGAGCTCAACCT |
| $Ilvleu-gfp_{uvr}$ | GAGCTCCCTATATTATGCTTTTCATTCATAAAAATGATTATCCATTGTTCAATCGTATCTAACTTTATA<br>TTTAACCTTTATATTGTAACAAATTTCAACTTAAATTTCTTATCTTTGAAACAGATTATCTATTCAAAG<br>TTAATTGTAAGAAAATTTAAAATATTTGTTGACATACTAAAGCAGATATAGTAAATTAAATTTATCAAA<br>TTTTTAGACAATTCTAACTATTAAAGTGATATATACCATTCACGGAAGGAGTATAATAAAATGCTTAAT<br>CAATATACTGAACATCAACCGACAACTTCAAATATTATTATTTTATTATACTCTTTAGGACTCGAACGT<br>TAGTAAATATTTACTAAACGCTTTAAGTCCTATTTCTGTTTGAATGGGACTTGTAAACGTCCCAATAAT<br>ATTGGGACGTTTTTTTATGTTTTATCTTTCAATTACTTATTTTTATTACTATAAAACATGATTAATCAT<br>TAAAATTT**ACGGGGG**AATTTACTatgtcaaaaggtgaagaattatttacaggtgtagtaccaattttag<br>tagaattagatggtgatgtaaatggtcataaattttcagtatcaggtgaaggtgaaggtgatgcaacat<br>atggtaaattaacattaaaatttatttgtacaacaggtaaattaccagtaccatggccaacattagtaa<br>caacatttacatatggtgtacaatgtttttcaagatatccagatcatatgaaaagacatgattttttta<br>aatcagcaatgccagaaggttatgtacaagaaagaacaatttcatttaaagatgatggtaattataaaa<br>caagagcagaagtaaaatttgaaggtgatacattagtaaatagaattgaattaaaaggtattgatttta<br>aagaagatggtaatattttaggtcataaattagaatataattataattcacataatgtatatattacag<br>cagataaacaaaaaaatggtattaaagcaaattttaaaattagacataatattgaagatggttcagtac<br>aattagcagatcattatcaacaaaatacaccaattggtgatggtccagtattattaccagataatcatt<br>atttatcaacacaatcagcattatcaaaagatccaaatgaaaaaagagatcatatggtattattagaat<br>ttgtaacagcagcaggtattacacatggtatggatgaattatataaataaTAATTGAGCTCAACCT |

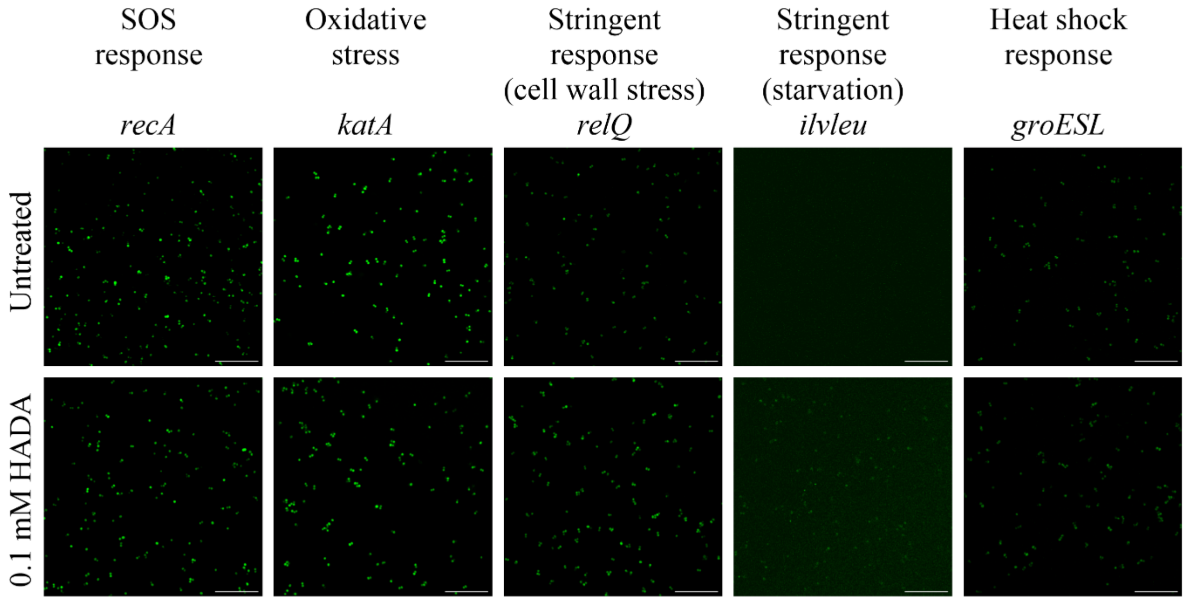


**Fig. S1 Incubation with HADA does not activate stress responses.** Reporter strains were grown to mid exponential phase with or without 0.1 mM HADA in TSB for approx. 2 h. Subsequently, samples were washed and resuspended in PBS before mounting on a glass slide with antifade. Samples were visualised using CLSM with a 488 nm laser to excite GFP. Image acquisition settings were optimised for each strain but kept constant across samples from different growth conditions. Scale bar: 20 µm.

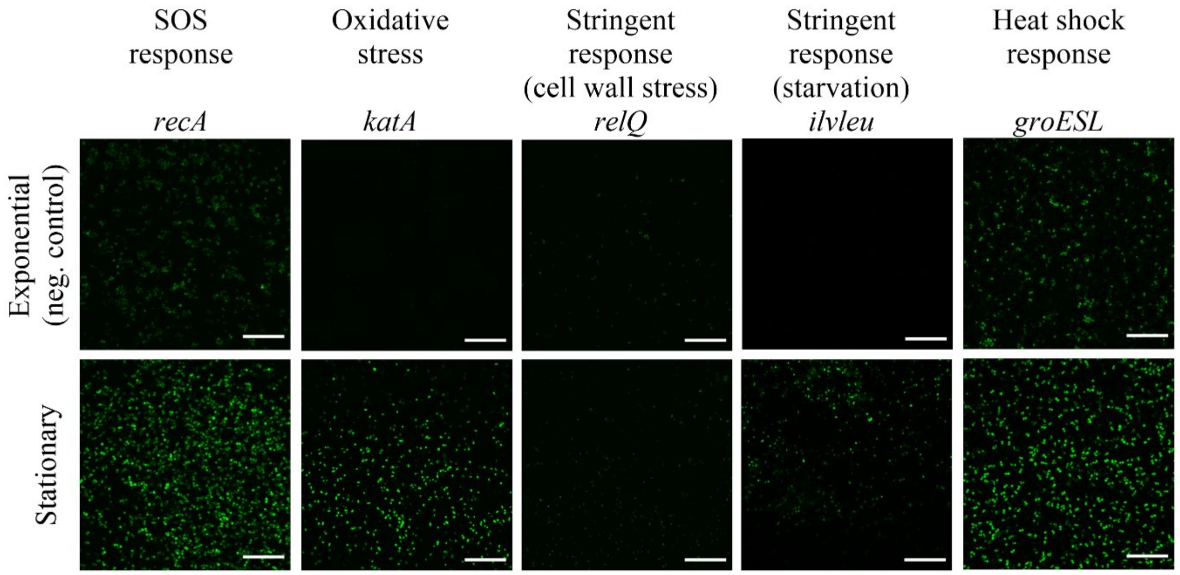


**Fig. S2 Most stress responses are activated in stationary phase cultures.** Reporter strains were grown to mid exponential phase with or without 0.1 mM HADA in TSB for approx. 2 h. Sationary phase cultures were grown overnight with 0.1 mM HADA in TSB. Subsequently, samples were washed and resuspended in PBS before mounting on a glass slide with antifade. Samples were visualised using CLSM with a 488 nm laser to excite GFP. HADA (data not shown) was used to locate the cells. Image acquisition settings were optimised for each strain but kept constant across samples from different growth conditions. Scale bar: 20 µm.

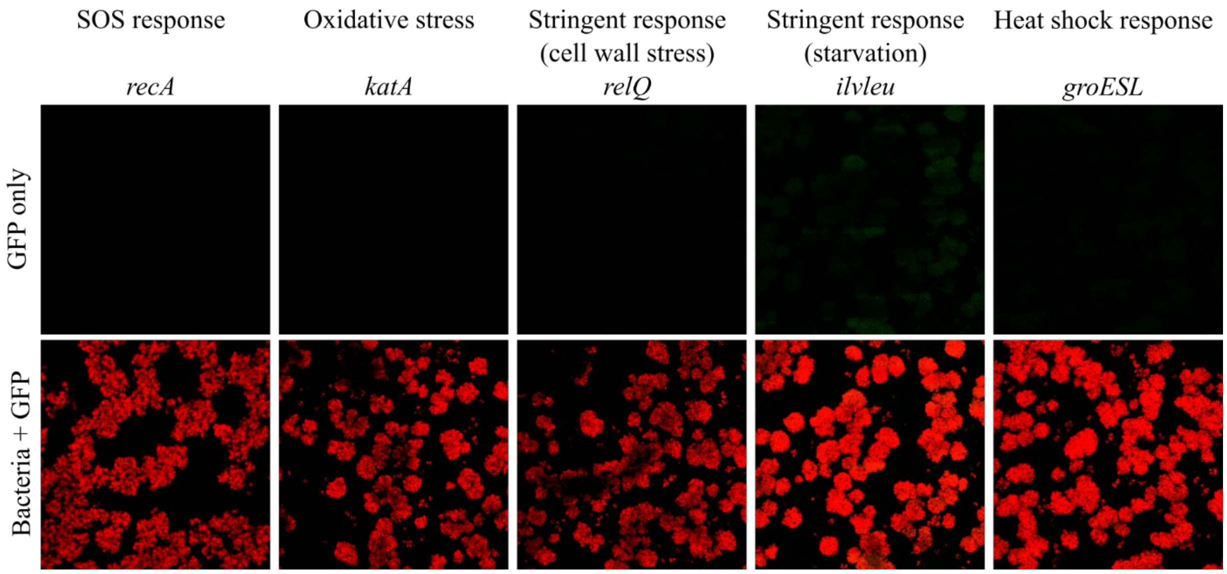


**Fig. S3. Empty vector control strain biofilm images obtained with the settings used for visualising each biofilm of the reporter strains.** *S. aureus* 29213 pKK30 was grown to mid exponential phase and resuspended in 100% serum with 0.1 mM HADA and inoculated in a well pre-conditioned with 100% human plasma. After 24 h growth, the biofilms were washed with PBS and imaged by CLSM. Green = GFP and red = HADA (peptidoglycan cell wall). n = 3 biological replicates. Scale bar: 20 µm.

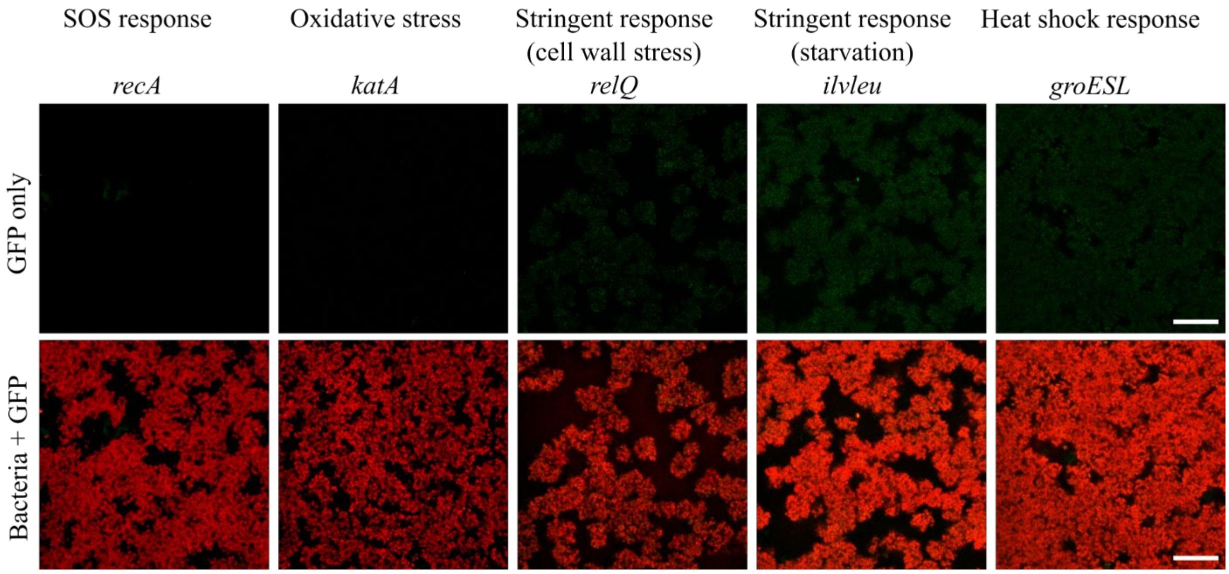


**Fig. S4. Activation of stress responses in biofilms grown in 10% serum diluted in TSB.** *S. aureus* reporter strains was grown to mid exponential phase and resuspended in 10% serum in TSB with 0.1 mM HADA and inoculated in a well pre-conditioned with 100% human plasma. After 24 h growth, the biofilms were washed with PBS and imaged by CLSM. Green = GFP and red = HADA (peptidoglycan cell wall). n = 3 biological replicates. Scale bar: 20 µm.

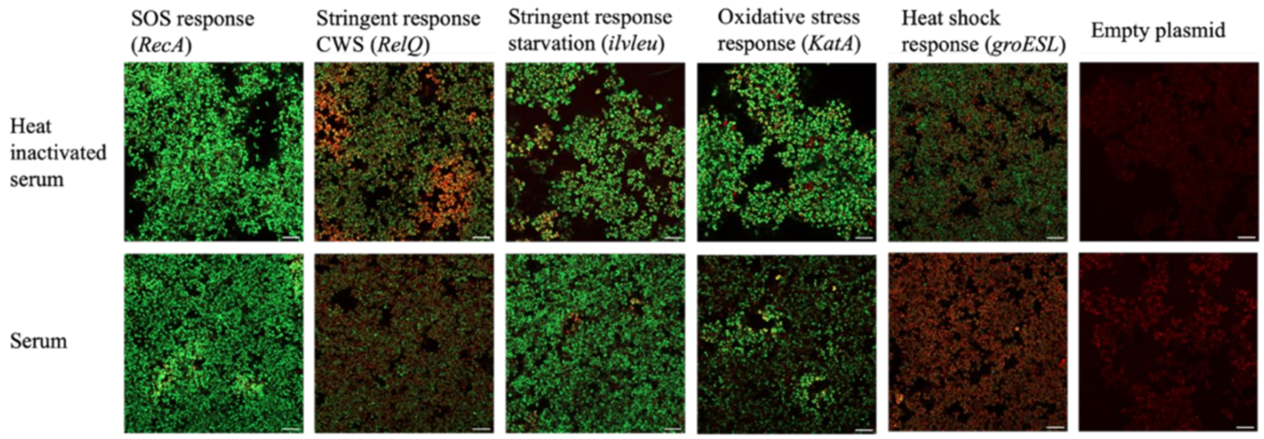


**Fig. S5. Activation of stress responses in biofilms grown in 100% serum or heat-inactivated serum.** *S. aureus* reporter strains were grown overnight and adjusted to OD600 = 0.1 and inoculated in a well pre-conditioned with 10% human-plasma for 1 h, before the liquid in the wells was aspirated and exchanged to either TSB, AUM, heat inactivated serum or serum. After 48 h growth, the biofilms were washed with PBS and imaged by CLSM. GFP fluorescence was visualized as a measure of activation of stress responses using a 401 nm laser to excite HADA and 488 nm laser to excite GFP. Images are 2D and acquired from approximately the bottom of the biofilm. Green = GFP, red = HADA (peptidoglycan cell wall). n = 3 biological replicates. Scale bar: 10 μm.
